# Delivering trait-enhanced varieties to African smallholders through a pangenomic breeding network

**DOI:** 10.1101/2025.08.07.667917

**Authors:** Fanna Maina, Jacques M. Faye, Alex W. Kena, Cyril Diatta, Moctar Issiakou Tankari, Ousseini Abdou Ardaly, Ousmane Seyni Diakite, Aissata Mamadou Ibrahim, Oumarou Abdoulye Moussa, Abdou Harou, Rabiou Abdou, Joseph Pascal Sene, Souleymane Bodian, Mbery Ndour, Diarietou Sambakhe, Bassirou Sine, Pana Kadanga, Eyanawa Akata, Nofou Ouedraogo, Benjamin Annor, Israel T. Tetteh, Muhammad Ahmad Yahaya, Abdoulaye Diallo, Ronald Kakeeto, Rachael K. Kisilu, Clarisse Pulchérie Kondombo, Tokuma Legesse, Lloyd Mbulwe, Joaquim Mutaliano, Emmanuel T. Mwenda, Gapili Naoura, Hortense Noëlle Apala Mafouasson, Kenneth Opare-Obuobi, Rekiya Otuchu Abdulmalik, Steven Runo, Tilal Sayed Abdelhalim, Louis Yalaukani, Joel Masanga, Chloee M. McLaughlin, Jesse R. Lasky, Avril M. Harder, Adam L. Healey, John T. Lovell, Portia Osei-Obeng, Sandeep R. Marla, Terry J. Felderhoff, Falalou Hamidou, Daniel Fonceka, Mohan Kumar Varma Chejerla, Baloua Nébié, Mark Nas Tamerlan, Amos Alakonya, Abhishek Rathore, Santosh Deshpande, Harish Gandhi, Linly Banda, Davina H. Rhodes, Clara Cruet-Burgos, Carl J. VanGessel, Geoffrey P. Morris

**Affiliations:** Institut National de la Recherche Agronomique du Niger, Niamey, Niger; Institut Sénégalais de Recherches Agricoles, Thiès, Sénégal; International Maize and Wheat Improvement Center (CIMMYT), Nairobi, Kenya; Department of Crop and Soil Sciences, Kwame Nkrumah University of Science and Technology, Kumasi, Ashanti, Ghana; Department of Soil & Crop Science, Colorado State University, Fort Collins, CO, USA; Université André Salifou, Zinder, Niger; West Africa Centre for Crop Improvement, University of Ghana, Accra, Ghana; Institut Togolais de Recherche Agronomique, Lomé, Togo; Institut de l’Environnement et de Recherches Agricoles, Ouagadougou, Burkina Faso; Institute for Agricultural Research, Samaru, Ahmadu Bello University, Zaria, Nigeria; Institut d’Economie Rurale, Centre Régional de Recherche Agronomique, Sotuba, Mali; National Agricultural Research Organization, National Semi-Arid Resources Research Institute, Soroti, Uganda; Kenya Agricultural and Livestock Research Organisation, Kenya; Ethiopian Institute of Agricultural Research, Melkassa Agricultural Research Center, Melkasa, Ethiopia; Zambia Agriculture Research Institute, Lusaka, Zambia; Instituto de Investigação Agrária de Moçambique, Ulónguè, Tete, Moçambique; Tanzania Agricultural Research Institute, Dodoma, Tanzania; Institut Tchadien de Recherche Agronomique pour le Développement, République du Tchad; Institute of Agricultural Research for Development, Yaoundé, Cameroon; Council for Scientific and Industrial Research-Savanna Agricultural Research Institute, Nyankpala/Tamale, Ghana; Department of Biochemistry, Microbiology, and Biotechnology, Kenyatta University, Nairobi, Kenya; Biotechnology and Biosafety Research Center, Agricultural Research Corporation, Khartoum North, Sudan; Department of Agricultural Research Services, Lilongwe, Malawi; Department of Biology, Pennsylvania State University, University Park, PA, USA; Genome Sequencing Center, HudsonAlpha Institute for Biotechnology, Huntsville, AL, USA; Department of Horticulture and Landscape Architecture, Colorado State University, Fort Collins, CO, USA; US Department of Energy Joint Genome Institute, Berkeley, CA, USA; Department of Agronomy, Kansas State University, Manhattan, KS, USA; International Crops Research Institute for the Semi-Arid Tropics – Sahelian Center, Niamey, Niger; Centre de coopération internationale en recherche agronomique pour le développement, Montpellier, France; International Crops Research Institute for the Semi-Arid Tropics, Patancheru, Telangana, India

## Abstract

Pangenomics has been promoted to accelerate breeding of orphan crops, but smallholder farmers in developing nations have seen little benefit so far. To address this gap, we built a global pangenomic breeding network, integrating African breeding programs, U.S. land grant universities, and international nonprofit research organizations. Here we demonstrate that pangenomics, when integrated with local crop improvement knowledge and global scientific partnerships, can facilitate breeding of drought and pest resilient varieties for smallholders. To breed trait-enhanced sorghum varieties with *lgs1*-*1* resistance to witchweed (*Striga hermonthica)* for smallholders in Niger, one of the world’s least developed nations, we used population genomics across local and global scales to develop *lgs1*-*1 Striga* resistance markers, and deployed them for rapid introgression of resistance into locally-preferred varieties. Genomic characterization, along with controlled experiments in laboratory, pot, field stations, and smallholder farms, confirmed *lgs1*-*1* resistance was introgressed without loss of essential local-preference traits. New pangenomic resources, including global resequencing and graph pangenomes, further accelerated design of broadly-applicable markers. Unlocking the potential of pangenomics for stress-resilience breeding depended on stakeholder input, strong inference, South-led decision support software, and a dense collaborative network. The experience of the network provides a scalable roadmap for collaborative pangenomic breeding of trait-enhanced varieties for the world’s lowest-resourced farmers.

---

Increasing the productivity and resilience of smallholder farming has been one of the most reliable and cost-effective ways of unlocking human flourishing over the past century^1^. Unfortunately, more than a billion people, mostly in sub-Saharan Africa, have yet to access benefits of improved agricultural production^1,2^. In response, major investments have been made in orphan crop genomics, but there has been little evidence of benefits to smallholders so far^3–5^. One approach to achieve the benefits of orphan crop genomics is empowering developing-country plant breeders to lead genomics-enabled crop improvement initiatives, as these scientists are best equipped to bridge local knowledge of smallholder communities with research capacity of the global scientific community^4,6,7^ (Fig. 1a). The pangenomes of orphan crops harbor abundant diversity for useful traits and signatures of selection to guide crop improvement. Successive improvements in surveys of orphan crop pangenomes^3,5^—from genotyping-by-sequencing (GBS; ∼0.1–1% of SNPs), to whole-genome resequencing (∼100% of SNPs), and multiple de novo reference genomes (SNPs and structural variants)—have further enhanced the potential value of bridging pangenomics and breeding in developing countries.

**Fig. 1.**
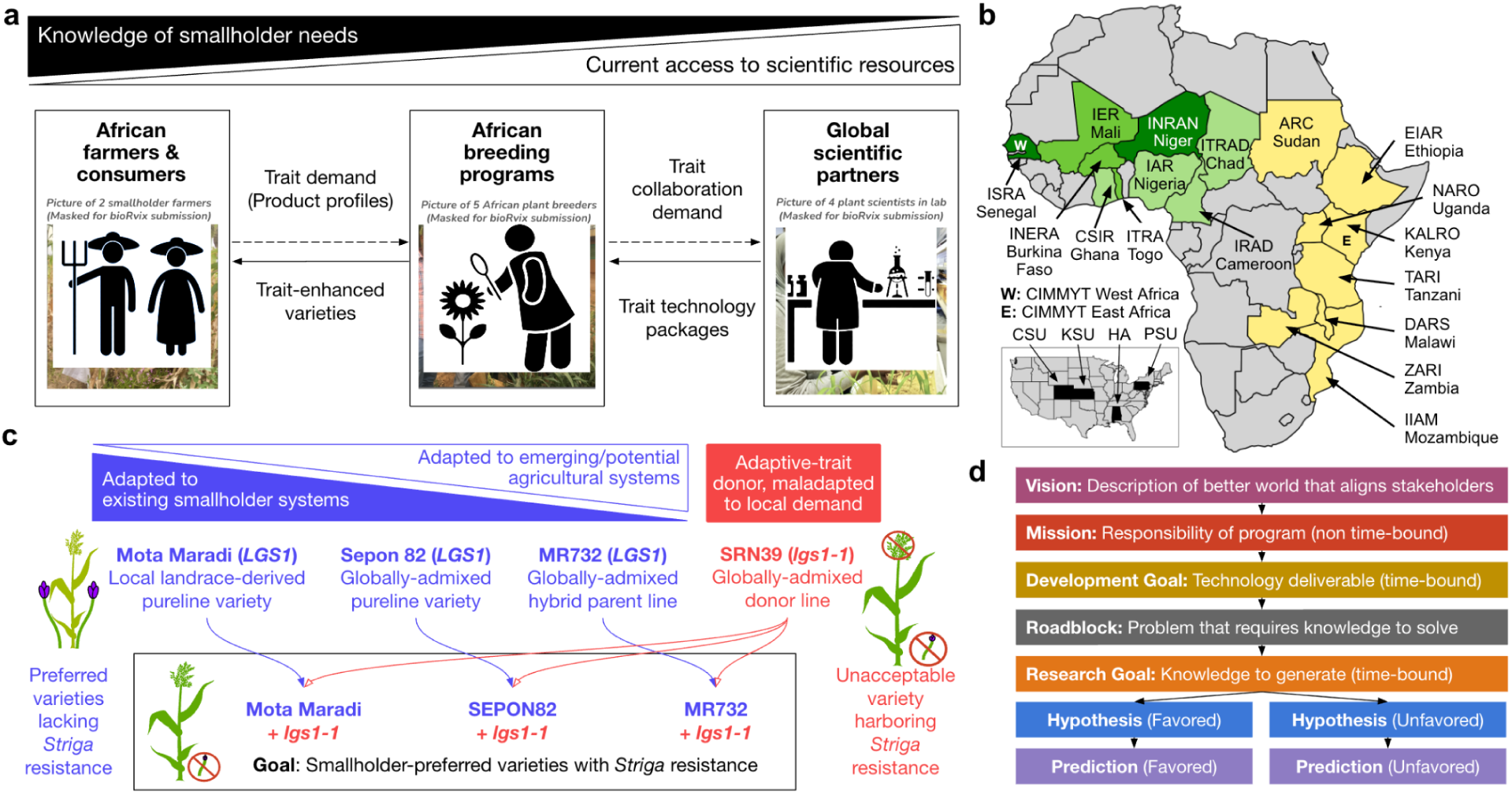
Design and implementation of a global pangenomic breeding network for African smallholder farmers. **a**, Framework for a global pangenomic breeding network that meets smallholder farmer needs. African smallholder farmers and farmer representatives (left; smallholders in a *Striga* infested sorghum field) provide trait demand to African breeding programs (center; African breeders at INRAN’s *Striga* resistance breeding nursery) who establish collaborations with global scientific partners (right; *Striga* resistance testing in a US partner laboratory); the collaborations lead to development of trait technology packages for breeding and, finally, to trait-enhanced varieties for African smallholders (leftward arrows). **b**, The global pangenomic breeding network we have developed for African sorghum improvement. The network links African NARI with limited facilities to other NARI with extensive facilities and global research organizations (inset). The network bridges the West (green shades) and East African (yellow) sorghum belts, with founding NARI hubs (dark green) and early scaling partners (medium green) noted. CIMMYT research hubs, which support the NARS network, in West (“W”) and East (“E”) Africa, and US partners, are noted (inset). **c**, Pangenomic breeding plan to introgress *lgs1-1 Striga* resistance into locally-preferred varieties from a variety that harbors resistance but was not adopted by smallholder farmers. The recipient varieties span a range of genetic backgrounds targeted to a range of agricultural systems. **d**, Formalized applied science framework adopted in the network, linking goal-setting logic chains and scientific method logic chains (extending the strong inference approach^32^; Extended Data Fig. 2). (Photo credits: G.P.M., G.P.M., F.M.)

National agricultural research systems (NARS) in African countries are anchored by breeding programs at national agricultural research institutes (NARI; Extended Data Table 1), and bolstered by partnerships with local universities, farmer organizations, local seed companies, and grain processors. NARI breeding programs face an exceptionally difficult task: improving desired traits (under directional selection; e.g. yield, pest resistance, drought tolerance) and recovering acquired traits (under stabilizing selection; e.g. flowering time, height, pigmentation), without established elite breeding populations^6,8–10^ and with few resources (e.g. 0.3–1.9 scientists per million tons of production in major sorghum producing countries)^11^. Failure to recover a single “must-have” trait, or to transfer the “winning trait” from a resistance donor, can cause rejection of new varieties by farmers^11^. We built our network around South-South (i.e. NARS-NARS) partnerships (Fig. 1b; Extended Data Fig. 1) to leverage complementary strengths among NARS. Contributors at U.S. land grant universities and nonprofit research organizations, including CGIAR centers (CIMMYT and ICRISAT), helped build network capacity (molecular breeding, pangenomics, strong inference) and supported development of trait technology packages (trait markers, adapted donor lines, and genotype-phenotype maps) (Fig. 1a,b; Extended Data Table 1). Outsourced marker genotyping platforms (such as Kompetitive Allele-Specific PCR; KASP) provide NARI breeders in the network the opportunity to deploy marker-assisted selection (MAS)^12,13^ using markers that distill pangenomic discoveries into tractable tools.

We sought to address the gap between pangenomic resources and smallholder needs for sorghum (*Sorghum bicolor* [L.] Moench), a multipurpose cereal crop (grain, forage, fiber) of smallholder farmers in Africa’s dry regions (semi-arid and subhumid savannah zones)^6,14^. A complex geographic mosaic of adaptation to climate, stressors, and cultures^15,16^ in this indigenous African crop provides abundant diversity for crop improvement, but hinders Green Revolution-style mega-varieties^11,14^. The top constraints on sorghum across Africa are drought^14^ and witchweed (*Striga hermonthica*), an obligate hemiparasite of sorghum and other cereals in Africa that causes up to 50% yield losses or field abandonment^17^. Thus, developing drought and *Striga* resilient varieties is a top breeding program priority across Africa^17,18^ (Supplementary Tables 1-4). *Striga* requires root-exuded host strigolactones to stimulate germination^19,20^ and in the 1950s low germination stimulant (LGS) resistance was discovered in a West African landrace (Framida)^19^. Unfortunately, limited agroclimatic adaptation and/or non-preferred end-use traits of existing *Striga*-resistant varieties has hindered adoption (e.g. <0.1% adoption of Framida and Framida-derivative SRN39)^11,21,22^. Deploying our African-led GBS mapping of genomic diversity^23–27^, and leveraging other groups’ trait discoveries (mapping of stay-green *Stg* drought tolerance quantitative trait loci [QTL]^28^ and cloning of the *lgs1 Striga* resistance gene^29^), we launched a pangenomic breeding network for African smallholders (Fig. 1b). We targeted drought and *Striga*-resilient sorghum as initial cases to build, test, and scale the network (Fig. 1c). To address lessons on product delivery^30^ and technology scaling^31^ failures in global development, we used a framework linking formal breeder goal-setting and strong inference^32^ scientific method (Fig. 1d; Extended Data Fig. 2). Here, we illustrate the approach for breeding of *Stg* drought tolerance (Extended Data Fig. 3) and *lgs1 Striga* resistance (detailed below).

## Developing markers using population genomics

*Striga* resistance can be conferred by large structural variants (*lgs1-1*; 34 kb, *lgs1-2*, 29 kb; *lgs1-3,* 30 kb) that delete the sulfotransferase gene *LGS1* (Sobic.005G213600)^29^ (Fig. 2a). A previously developed *lgs1* PCR marker (for *lgs1-1* in SRN39) discriminates resistant and susceptible alleles in the INRAN-Niger breeding program, but was not tractable for MAS in Niger due to a lack of equipment and reliable power (Extended Data Fig. 1; at network launch Niger was ranked 188th of 188 on UN Human Development Index)^33^. We reasoned we could develop an outsourced KASP marker for *lgs1*, suitable for MAS at INRAN and other programs in the region, by identifying GBS SNPs in linkage disequilibrium (LD) with *lgs1* deletions. Lines carrying an *lgs1* deletion allele should have mostly missing data (80–100%) in the *lgs1* deletion region (based on 21 GBS SNPs in the interval), while *LGS1* genotypes should not. GBS-based inference suggested a high frequency of deletion alleles (e.g. 7% and 9% for initial focal countries of Niger and Senegal, respectively) and included known resistant genotypes, however missing SNPs may be due to low coverage (an inherent limitation of GBS^34^) rather than *lgs1* deletion (permutation test: 3% false positive rate; Fig. 2b). Global germplasm (*N* = 1724)^29^ inferred to have large deletions indicated 224 African landraces are possible *lgs1* carriers (Fig. 2c).

**Fig. 2.**
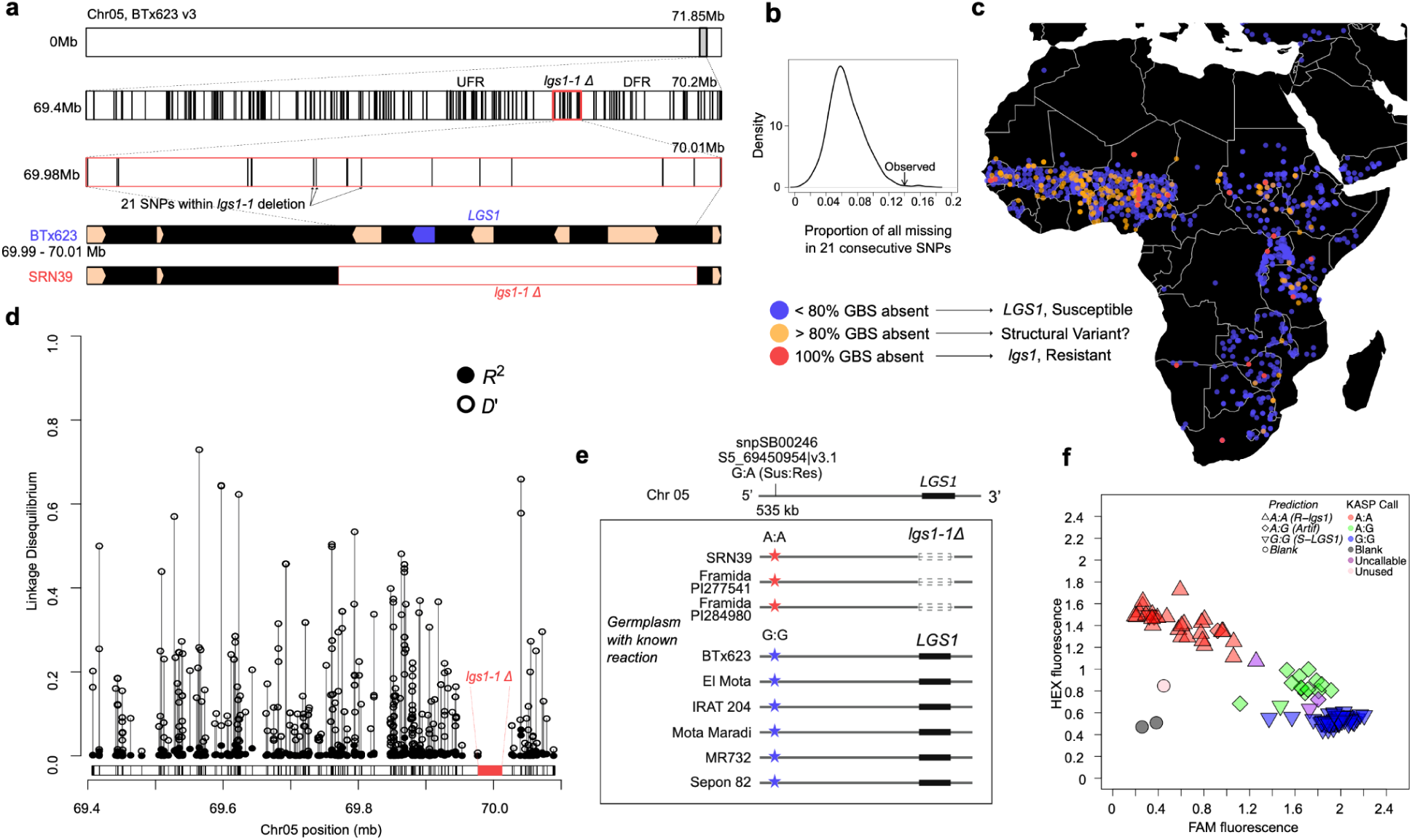
Population genomics analyses to develop an *lgs1* marker to transfer *Striga* resistance. **a**, Genomic characterization of *LGS1* region on Chr05 using genotyping-by-sequencing (GBS) SNPs. *LGS1* region (69.40–70.02 Mb|v3.1) includes the *lgs1-1* deletion region in SRN39, encompassing 5 genes including Sobic.005G213600 (*LGS1* gene), as well as the upstream (UFR, 69.40–69.97 Mb|v3.1) and downstream (DFR, 70.01–70.02 Mb|v3.1) flanking regions. **b**, Permutation test to estimate frequency of false positives using GBS-based inference of *lgs1* deletion (21 missing GBS SNPs). **c,** Geographic origin of georeferenced accessions in Africa and West Africa color-coded based on the hypothesized *LGS1* wild-type (blue) and *lgs1* deletion (red, validated with known resistant genotypes SRN39: PI656027, 54.K.94: PI533752, and Framiola: PI533976). Deletion of *LGS1* was inferred based on the number of missing GBS SNPs within the “*LGS1* region” (69.97–70.011 Mb). **d**, One-dimensional linkage disequilibrium analysis (*R*^2^ and *D*’) for GNS-inferred *lgs1* allele versus SNP alleles in the extended *LGS1* region (*lgs1-1* deletion, UFR, DFR); orange stars indicate the position of single nucleotide polymorphisms (SNPs) tested for marker development. **e**, Identification of a SNP with perfect LD for the INRAN breeding program. **f**, KASP genotyping result for *lgs1-1* marker (snpSB00246) on *Striga* susceptible (BTx623, El Mota, IRAT204, Mota Maradi, MR732) and resistant (SRN39) genotypes. Expected KASP calls for each sample are represented by upward triangles (predicted A:A), downward triangles (predicted G:G), diamonds (predicted A:G), and circles (Unknown or no *a priori* information) are *a priori* hypotheses based on known *Striga* resistance or susceptibility.

To identify GBS SNPs in LD with *lgs1* deletions, suitable for development of KASP markers, we calculated *R*^2^ and *D*’ for SNPs (*N* = 342) in the genomic regions flanking the *lgsl-1* deletions (UFR: 577 kb, DFR: 100 kb) across global sorghum germplasm (*N* = 2,599) (Fig. 2d, Extended Data Fig. 4). The *D*’ ranged widely but was high for some SNPs (0–0.73), indicating some SNPs could be locally predictive of *lgs1* deletions. However, *R*^2^ of SNPs versus the GBS-inferred *lgs1* deletion were all low (0–0.06), suggesting that no SNP can be converted into a globally-predictive *lgs1* marker. We advanced 10 SNPs with contrasting alleles for parental genotypes to KASP marker conversion at our outsourced genotyping provider (Intertek Agritech, Sweden), and found eight markers that passed ‘SNP Quality Assessment’ (Supplementary Table 5). Finally, we advanced a single marker (snpSB00246; S5_69450954|v3.1), assaying a SNP at 535 kb upstream of *lgs1* that was in perfect LD (*R*^2^ = *D*’ = 1) for resistant (*N* = 4) versus susceptible genotypes (*N* = 9) within the INRAN program (Fig. 2e, Extended Data Fig. 4). Due to the low *R*^2^ of the target SNP with the GBS-based *lgs1* deletion calls, this marker would not be expected to be trait-predictive in other global breeding programs. However, based on the high *D*’ among INRAN’s parent lines, this marker should be locally trait-predictive and suitable to advance a marker-assisted backcross (MABC) program.

Testing the KASP marker, we found genotyping calls for snpSB00246 were consistent with the expected allele for known susceptible and resistant genotypes (Fig. 2f). Further, for the artificial heterozygote samples, almost all samples (11/12) were called correctly as A:G heterozygotes, validating the marker for selection of heterozygotes during MABC. We validated the markers for MAS in two biparental breeding families, observing consistent SNP calling across technical replicates of parents and F_3_ progenies (Supplementary Table 6). Note, given that weak clustering and some erroneous genotype calls were observed with snpSB00246, we later designed a reverse strand marker (snpSB00487, at the same SNP), which has better genotypic discrimination (Extended Data Fig. 5). Further, we developed decision support software (panGenomeBreedr)^35^ designed by developing-country molecular breeders (A.W.K. and others) to facilitate marker design and MAS. The functions incorporate strong inference principles for marker development and deployment, using color- and symbol-coding based on a priori hypotheses and predictions to highlight discrepancies in controls (i.e. known homozygotes of each class, artificial heterozygotes) (Fig. 2f, Extended Data Fig. 6). Therefore, panGenomeBreedr quickly and clearly identifies failures in marker development, genotyping processes, or germplasm management, to drive improvement of breeding operations.

## Rapid introgression of *lgs1-1* resistance

To rapidly breed *Striga*-resistant sorghum varieties for smallholders, we launched MABC programs using three locally-preferred varieties as recurrent parents (Fig. 3a and Extended Data Fig. 7). To transfer *Striga* resistance, we used the *lgs1-1* KASP marker (snpSB00246 and snpSB00487) for several generations of MAS, as well as marker-based quality control during successive generations of line fixation. In an effort to rapidly test our favored hypothesis of successful marker-assisted trait introgression, we (F.M.) first used a laboratory germination assay at a *Striga* quarantine facility in the U.S. (Fig. 3b, Upper inset: positive control, synthetic strigolactone GR24; Lower inset; DI water, negative control). As expected, root exudates of the susceptible parents (Mota Maradi, MR732, and SEPON82) induced *Striga* germination, while exudates of the resistant donor (SRN39) did not (Fig. 3b). Importantly, the *lgs1-1* introgression lines (ILs) (Lines 7, 10, 11, 13, 15, 17, 19) induced almost no *Striga* germination (Fig. 3b) (*P* > 0.01). To deploy strong inference within the breeding program, we had also retained sibling ILs carrying the *LGS1* susceptible alleles for use as negative controls. As expected, almost all the *LGS1* sibling ILs (Lines 14, 16, 18, 9, 12) induced *Striga* germination at the same rate (62–71%; *P* = 0.3) as the recurrent parents (though surprisingly two of the sibling ILs, Lines 6 and 8, did not induce *Striga* germination, either due to experimental noise, uncoupling of the marker and causative variant, or introgression of uncharacterized resistance alleles from SRN39).

**Fig. 3.**
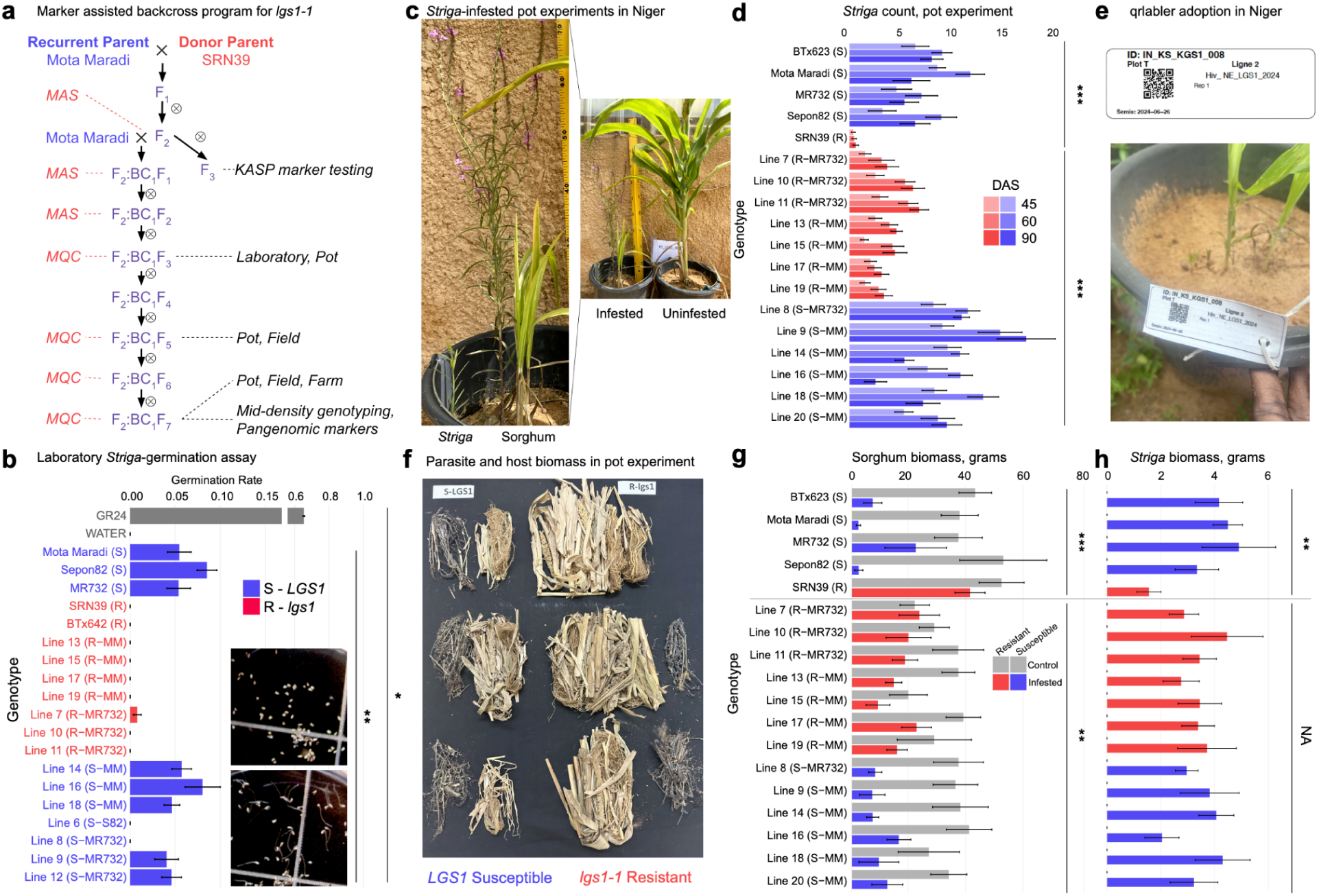
Effective introgression of *Striga* resistance into locally-preferred genetic backgrounds with *lgs1* marker. **a**, Marker-assisted backcross scheme to breed *Striga*-resistant Mota Maradi via *lgs1-1* introgression from SRN39. Steps of marker-assisted selection (MAS) and marker-based quality control (MQC) with *lgs1* KASP markers (snpSB00246 or snpSB00487) are noted in red text. **b**, Germination rate for progenies (BC_3_F_3_ and F_2_:BC_1_:F_3_) including inbred lines. Genotypic classes are indicated in *LGS1* (blue) representing G:G homozygote class and in *lgs1* (red) representing A:A homozygote class. Inset images of preconditioned *Striga* seeds 72 hours after exposure to GR24 (top) and water (bottom). **c**, *Striga* resistance pot experiment in Niger using artificial infestation of *Striga* from Niger. Host plant comparison in infested versus uninfested control (*Striga*-free) pot. **d**, *Striga* counts (45, 60, and 90 days after sowing) for Mota Maradi ILs, MR732 ILs, and parent lines in the pot assay. **e**, Use of qrlabelr RShiny app for barcode labeling in the pot experiment is an example of South-South capacity building. **f**, Dry biomass of sorghum (yellow material; inner columns) and *Striga* (gray material; outer columns) for representative *LGS1* ILs (left) and *lgs1-1* ILs (right). **g**, Sorghum biomass (above- and below-ground) and **h**, *Striga* biomass (above ground) for uninfested control pots (gray) versus *Striga*-infested pots (blue: *lgs1* lines; red: *LGS1* lines). (Photo credits: F.M.)\

*Striga* infestation may be affected by soil and climate factors, and *Striga* itself harbors ecotypic variation in response to host strigolactone profiles, and effectiveness of *lgs1* varies by *Striga* ecotype^36,37^. Following strong inference, we considered the unfavored hypothesis that *lgs1-1* resistance is ineffective in Niger. Over 3 years of trials, we tested the effectiveness of *lgs1-1* introgression in pot assays in Niger, using *Striga* from Nigerien smallholder farms and local soil (Fig. 3c). A substantial reduction of the number of emerged *Striga* plants (62–71%, *P* < 0.001) was observed for *lgs1-1* ILs compared to their *LGS1* sibling ILs (Fig. 3d). Interestingly, however, *lgs1-1* ILs had substantially higher *Striga* counts than donor parent SRN39 (90 days-after-sowing mean of 4.6 vs. 0.6), suggesting that SRN39 may harbor additional resistance QTL, not identified in the original mapping studies^38^, that are expressed in this environment. We also note, as a further example of South-South capacity-building, that as the scale of trials increased (Extended Data Table 2), the INRAN breeding program (F.M. and staffers) adopted qrlabelr barcode-labeling RShiny app^39^ (Fig. 3e). This app was designed by an African NARS scientist (A.W.K.) to help NARS implement foundational principles of data management (e.g. unique identifiers, standardization, traceability) and adopt open-source digitized breeding tools (e.g. BrAPI-compliant^40^ software including digital field books^41^ and breeding management systems^42^).

The reduction in *Striga* counts is promising, but not sufficient to conclude that *lgs1-1* improves aspects of performance that are relevant to smallholder farmers. Encouragingly, we further observed substantially greater sorghum biomass (above and below ground) for *lgs1-1* ILs compared to *LGS1* sibling ILs (Fig. 3f; yellow plant material, Fig. 3g), directly establishing a benefit of *lgs1-1* for forage yield. Notably, the effect was observed only in *Striga*-infested pots, not control pots (gray bars), establishing that *lgs1-1* confers *Striga* resistance per se (genotype-environment interaction). In striking contrast, we observed no difference in *Striga* biomass (Fig. 3h, gray biomass in Fig. 3g) between *lgs1-1* ILs and *LGS1* sibling ILs. This suggests that *Striga* ecotypes that are able to germinate on *lgs1-1* genotypes perform well afterwards, raising the worrying possibility that large-scale *lgs1-1* deployment could lead to rapid evolution of *Striga* that circumvent *lgs1*, unless additional resistance genes are pyramided onto the *lgs1* background^43^. Interestingly, there is further evidence that SRN39 harbors additional (non *lgs1*) *Striga* resistance alleles, as there was significantly less (58%; *P* < 0.01) *Striga* biomass in SRN39 pots compared to susceptible parent lines or the ILs. Together, these controlled-environment experiments established the effectiveness of *lgs1-1* MABC for Niger, and provided a basis to advance the *lgs1-1* ILs towards varietal release via multi-environment field trials.

## Varietal development from genome to farm

Adoption of new crop varieties with desired traits often fails without recovery of background acquired traits, such as agronomic adaptation or end-use acceptability^44^. The use of foreground selection only in the early stages of MABC (due to a paucity of background KASP markers and decision support tools at the time) left open two major avenues for failure: recombination between the marker SNP and the *lgs1-1* deletion (i.e. failed foreground selection for desired trait) and/or unacceptable alleles from the donor parent due to Mendelian segregation (i.e. failed recovery of acquired traits). We first characterized introgressions genome-wide in the ILs using a new mid-density genotyping assay (DArTag), designed to include highly-informative background markers and trait-predictive markers (see Methods), and panGenomeBreedr software (Fig. 4a). Introgressions of the SRN39 *lgs1*-*1* haplotype were successful with the marker allele and deletion remaining in linkage phase (Extended Data Fig. 8); future efforts using flanking markers and large families could recover more precise introgressions, while avoiding additional backcross generations that would delay varietal release. We assessed recurrent parent genome recovery at 16 loci that condition variation for traits classified in the regional market segments as must-have “basic traits” (*Tan, Ma,* and *Dw* genes) or “value-added traits” (*Stg* and *bmr* genes) (Supplementary Tables 1-4). Overall, recurrent parent haplotypes were observed at most loci (70% and 81% for Mota Maradi and MR732, respectively), suggesting that local preference traits had been recovered. Detailed examination of causative variant haplotypes for key traits provides insights on how pangenomic breeding could be optimized as more NARIs implement MAS and the number of known causative variants increases (Extended Data Fig. 8).

**Fig. 4.**
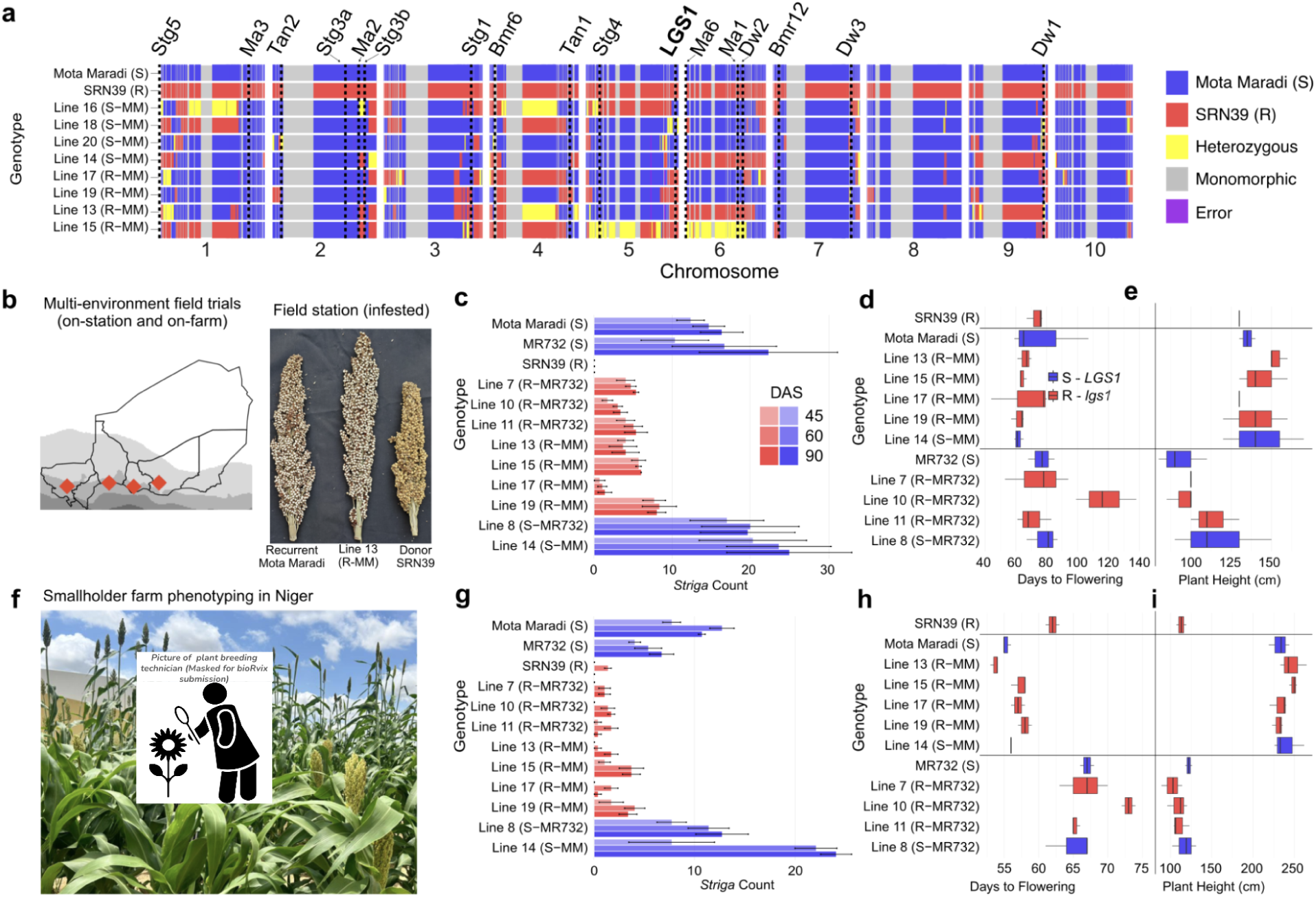
Precise introgression of *Striga* resistance and recovery of farmer-preferred traits. **a**, Genome-wide haplotypes for parent lines and marker-assisted backcross (MABC) progenies (F_2_:BC_1_F_7_) from the Mota Maradi MABC program. The haplotypes were analysed with a mid-density genotyping assay (DArTag) and visualized in panGenomeBreedr, noting *LGS1* and 16 genes/QTL that condition other traits that are specified in the target product profiles (Supplementary Table 2). **b**, Field phenotyping locations across Niger (color scale represents precipitation gradient) and panicles of parent and progeny lines collected on-station, showing recovery of recurrent parent phenotype in the ILs. **c**, Field phenotyping of *Striga* emergence (DAS: days after sowing) in controlled-infestation on-station trial in Niger (Konni) for Mota Maradi (MM) and MR732 ILs (BC_3_F_7_ and F_2_:BC_1_:F_7_) and parent lines, as a part of a set of multi-environment trials. **d**, Flowering time and, **e**, plant height phenotypes in the same experiment. **f**, Field phenotyping of *Striga* emergence, **g**, in a naturally-infested smallholder farm in Niger (Birni-N’Konni) for the same lines. **h**, Flowering time and, **i**, plant height phenotypes in the smallholder farm experiment. Height differences between recurrent parents represent traditional-style (tall: Mota Maradi) or Green Revolution-style (medium: MR732). (Photo credits: F.M.)

Adaptive traits that are promising in controlled environments (laboratory or pot experiments) can fail under agronomically-relevant field conditions^45,46^ (Fig. 4b). On-station field trials under controlled *Striga* infestation in Niger showed that *lgs1-1* introgression confers resistance in both Mota Maradi and MR732 backgrounds, with 79% reduction of *Striga* counts in *lgs1-1* ILs compared to *LGS1* sibling ILs (*P* < 0.001) (Fig. 4c). By contrast, sibling *LGS1* ILs were heavily infested (similar to the recurrent parents, *P* = 0.3), confirming that improved ILs performance is due to *lgs1* per se. Simultaneous to *Striga* phenotyping, on-station field trials determined the MABC scheme sufficiently recovered locally-preferred background traits. Flowering time is the most important local adaptation trait for cereals in regions with precipitation seasonality^14^. While the trait donor, SRN39, has similar flowering time to the recurrent parents, Mota Maradi and MR732, cryptic genetic heterogeneity between donor and recurrent parents could create transgressive segregation and result in failed adoption. On-station trials showed that *lgs1* ILs recovered optimal flowering time (∼70 days), other than one (Line 10) that was unacceptably late (∼110 days) (Fig. 4d). Plant height is another trait that is essential for smallholder farmer acceptance, as a determinant of secondary (forage and building material) yield characteristics. The ILs all recovered acceptable plant height, matching the recurrent parent (Fig. 4e).

Limited adoption of modern varieties by smallholder farmers^44^ has been attributed in part to varietal testing in unrepresentative on-station trials (e.g. under favorable fertilization, irrigation, and pest management) rather than representative on-farm trials (e.g. under limited nutrient, water limitation, and pest pressure)^47^. From a genetics standpoint, it is critical to exclude the hypothesis of unfavorable genotype-environment interaction between the on-station environment and the on-farm target population of environments^14^. From a behavioral and economics perspective, ensuring similarity of improved varieties with locally-preferred varieties should also facilitate adoption. Thus, we tested *Striga* resistance and background trait recovery on smallholder farms near Birni-N’Konni, and near Zinder, over 400 km from Birni-N’Konni and 740 km from Niamey (Fig. 4f). Under natural *Striga* infestation in a smallholder’s field near Birni-N’Konni, *lgs1-1* introgression reduced *Striga* counts by 88% compared to recurrent parents and *LGS1* sibling ILs (ANOVA *P* < 0.001; Fig. 4g), with all *lgs1-1* ILs performing as well or better than they did on-station (Fig. 4c). Similarly, in smallholder fields near Birni-N’Konni (*Striga*-infested) and Zinder (uninfested), phenotyping of flowering time (Fig. 4h), plant height (Fig. 4i), and grain yield components (Extended Data Fig. 9) demonstrate the *lgs1-1* ILs (other than Line 10, which was unacceptably late, as in on-station trials) recovered agronomic traits that are essential for varietal release and adoption.

## Scaling pangenomic breeding across Africa

We initially scaled the pangenomic breeding network from hubs at ISRA-Senegal and INRAN-Niger to early-adopter programs, INERA-Burkina Faso (*Stg* MABC; Extended Data Fig. 3) and ITRA-Togo (*lgs1* MABC; Extended Data Fig. 7), in the West African subregion (Fig. 1b). Based on promising findings in these countries (Extended Data. Fig. 3, Figs. 2–4), we sought to expand the approach across almost all the NARI in the East and West African sorghum belts, as a part of the African Dryland Crop Improvement Network (ADCIN) (Fig. 5a; shaded regions). The countries represented in ADCIN account for 89% of African sorghum production area and quantity (and 62% and 40% global production area and quantity)^48^. Within ADCIN, the NARI and CIMMYT have formalized target product profiles that capture demand from local stakeholders (smallholder farmers, consumers, processors, seed enterprises), and developed prioritized regional market segments with a distributed NARS leadership structure (Supplementary Tables 1-4). Genotyping of the GBS-based *lgs1* markers (snpSB00246/snpSB00487) across ADCIN showed high prevalence of *lgs1-*associated alleles (30% and 40% of West and East African germplasm, respectively). However, these are likely to mostly represent false positives for *lgs1*, since the marker allele has a high MAF, and global LD between marker and *lgs1* was low (Fig. 2c, 2d). Therefore, we used new pangenome resources, including whole-genome sequencing (WGS; *N* = 2,144) and de novo reference genomes (*N* = 33) capturing global diversity of georeferenced landraces and key improved varieties^49^, to characterize limitations of the first generation markers and design new markers that are scalable across Africa (Fig. 5b).

**Fig. 5.**
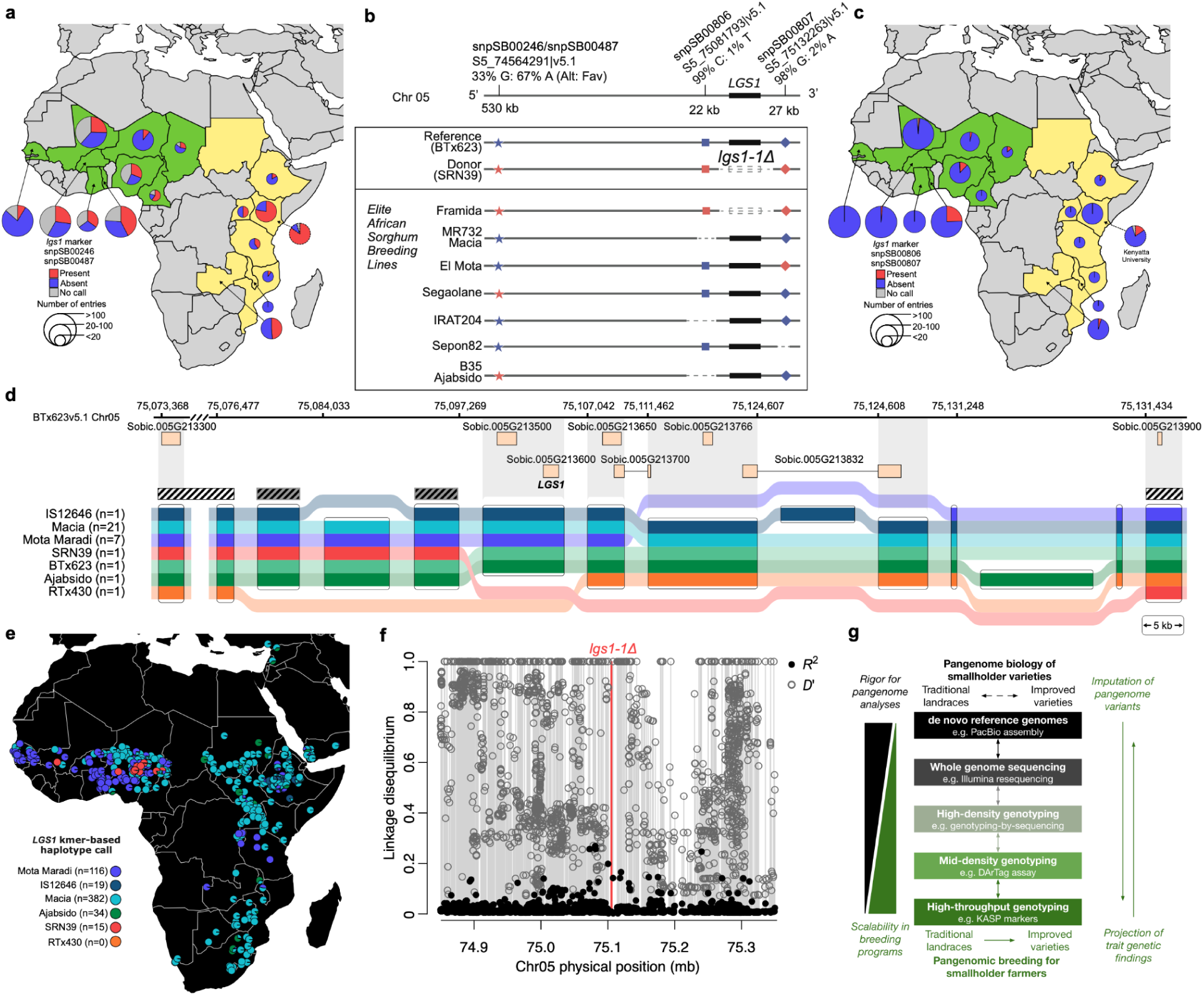
Using pangenomic resources to design markers that are effective from local to global scales. **a**, Continent-wide scaling of outsourced KASP genotyping across the African Dryland Cereals Improvement Network (ADCIN; 17 African NARI breeding programs, one African trait discovery program) demonstrates the feasibility of this MAS approach, but reveals limitations (high-false positive rate) of the GBS-based KASP markers (snpSB00246, snpSB00487) at a global scale. **b**, Whole-genome resequencing resources were used to characterize the limitations of the first generation markers (snpSB00246, snpSB00487) and develop improved markers (snpSB00806, snpSB00807). **c**, Improved pangenome-based KASP markers are more reliable in the global network, with high inferred *lgs1* frequency found only in expected programs (region of origin: IAR-Nigeria and ITRAD-Chad; programs selecting *lgs1*: ITRA-Togo and Kenyatta University). **d**, Tubemap graph of 33 pangenome references highlighting structural complexity of *LGS1* locus. Structural variation leads to causative *lgs1* resistance alleles and defines optimal regions for design of global markers (hatched boxes: black, considering all haplotypes; gray, ignoring the Tx430-like haplotype not observed in Africa). **e**, kmer-based inference of *lgs1* status in georeferenced African landrace accessions (color-coded as in panel c) contrasting resistant *lgs1-1* (red) genotypes versus susceptible *LGS1* genotypes (cool shades). **f**, Linkage disequilibrium analysis of kmer calls of *lgs1* versus resequencing SNPs for kmer-based design of improved *lgs1* resistance markers. While no nearby SNP has high *R*^2^ (≫50%) with *lgs1-1*, many nearby SNPs have high *D*’ (∼1), demonstrating that combinations of KASP markers could be used to accurately infer *lgs1-1* presence using decision support tools. **g**, Integrated pangenomic technology links high-resolution pangenomic discoveries (black) on smallholder landraces to low-cost selection tools for breeders (green) via decision support software and various scales of genomic/genotyping tools to breed trait-enhanced varieties for smallholders (rightward arrow).

LD analysis with a 1,676 WGS panel showed no association (*R*^2^ =0.001) at a global scale between WGS-inferred *lgs1-1* and the snpSB00487 SNP (S5_74564291|v5.1; note we transition to the improved v5.1 reference^49^ at this point). Further, resequencing data at S5_74564291|v5.1 (67% *lgs1*-associated A allele) indicated *LGS1-*containing B35 (BTx642), Ajabsido (Sudan), Segaolane (Botswana), and Marupantse (Botswana) will be incorrectly called as *lgs1*, since they harbor the *lgs1*-associated A allele (Fig. 5b). Thus, we selected two flanking SNPs tightly linked with *lgs1-1* (S5_75081793|v5.1 and S5_75132263|v5.1) for pangenome-based marker development (Fig. 5b, Extended Data Fig. 10). Genotyping the INRAN program breeding lines for S5_75081793|v5.1 (snpSB00806) and S5_75132263|v5.1 (snpSB00807) showed three distinct clusters, homozygous *lgs1-1*, homozygous *LGS1*, and heterozygous (Extended Data Fig. 10). Genotyping with snpSB00806 (MAF = 1%) suggested SRN39 contained *lgs1-1*, while BTx623, El Mota, and Mota Maradi contained *LGS1*. Genotyping of *LGS1*-containing MR732 with snpSB00806 would erroneously suggest it harbors *lgs1-1*, since it is clustered with empty controls. However, this can be explained by the pangenome data, which revealed that a flanking deletion covering S5_75081793|v5.1 in MR732 resulted in the loss of the primer annealing site and the clustering with empty controls (Extended Data Fig. 10). Given that SRN39, El Mota, and Mota Maradi share the same allele at S5_75132263|v5.1 (MAF = 2%), snpSB00807 would incorrectly call the presence of *lgs1-1* in El Mota and Mota Maradi, but it accurately calls *LGS1* in BTx623, MR732, IRAT204, and Macia. Based on these genotyping results, we are using snpSB00806 to introduce *lgs1-1* from SRN39 into El Mota and Mota Maradi, and snpSB00807 to introduce *lgs1-1* from SRN39 into IRAT204 and Macia, major adopted varieties for East and West Africa, respectively^11,50^. Thus, even in a crop with a relatively stable genome such as sorghum (e.g. compared to maize^51^), pangenomic maps of structural variants, along with pangenomic decision support tools, will be essential to guide marker design and deployment across global breeding networks.

Finally, to leverage graph pangenome approaches for marker development, we characterized the *LGS1* locus in the pangenome reference sequences, which harbors several derived variants that cause major *Striga* resistance (Fig. 5c), and projected them onto resequenced global georeferenced accessions (Fig. 5d). Several major structural variants on or around *LGS1* are present among our reference genomes (Fig. 5c) including the rare 28 kb deletion typical of *lgs1-1* resistant genotypes like SRN39 (red haplotype), a high-frequency (7/33 references; 21%) 14 kb deletion downstream of the *LGS1* gene that interferes with flanking markers (blue; Mota Maradi), and a 29 kb deletion known in *lgs1* resistant line RTx430 (orange)^52^, with breakpoints that correspond to the *lgs1-2* allele^29^. The pangenome reference also revealed previously unknown alleles, including two large insertions downstream of *LGS1* (13 kb, dark green; 8.9 kb, navy). Notably, these large structural variants must be avoided in KASP marker development to maximize utility of markers across populations (Fig. 5d; hatched boxes). To track these variants across global sorghum diversity, we counted exact matches to allele-specific kmer-dictionaries^49^ and calculated ancestry probabilities for all six major structural alleles and 2,144 resequenced genotypes (Fig. 5e). The kmer analysis locates the origin of the *lgs1-1* allele in landraces of the central Sahel (present-day northern Nigeria; red pie charts) from which it moved (via Framida, a Nigerian/Chadian landrace) to SRN39 in Sudan^29^, and on to the global breeding community. Notably, *LGS1* proximal variants were in low LD with the SRN39 haplotype (*R*^2^ < 0.3, *D*’ ≤ 1) (Fig. 5f), highlighting the need for multiple markers and decision support to track multiple causative alleles across global breeding networks.

## Conclusions

Our next step for delivery from the pangenomic breeding network is the release of *lgs1*- and *Stg-*enhanced varieties to the regional seed system^50^. Based on favorable evidence of *Striga* resistance (Fig. 2–4) and recovery of must-have local-preference traits (Fig. 4) the INRAN breeding program is advancing trait-enhanced varieties for release in Niger and nearby countries via harmonized regional seed regulation. Development of an *lgs1*-enhanced hybrid would take additional steps. Since MR732 is a pollinator parent for the NAD-1 hybrid variety (not a variety itself), *lgs1-1* is recessive, and the NAD-1 seed parent (ATx623) carries the dominant *LGS1* susceptible allele (Fig. 3f, 5c), development of an *lgs1-1* hybrid variety would require marker-assisted introgression into the maintainer BTx623, followed by backcross conversion of *lgs1*-BTx623 to a male-sterile *lgs1*-ATx623 seed parent. However, pangenomics findings on the prevalence of *lgs1-1* resistance (Fig. 5c) suggest an alternative “prebreeding by sequencing” approach: using the pangenomic database to identify existing seed parents (A/B lines) that harbor *lgs1* resistance and fast-tracking hybrids of these A-lines with MR732+*lgs1-1* ILs to performance trials. Deployment of *lgs1-1* varieties is unlikely to eliminate *Striga* constraints, especially since *Striga* biomass, and presumably seed production, seems to be unaffected by *lgs1-1* (Fig. 3h). A durable strategy may require pyramding traits^21,53,54^, deploying suicidal germination to reduce the parasite seed bank^55^, and scaling management practices that reduce infestation.

Beyond the particular trait-enhanced varieties we developed, we demonstrated the feasibility and utility of a global pangenomic breeding network approach (Fig. 5g), which could be scaled to other staple smallholder crops (e.g. maize, millet, cowpea, common bean) that underlie food security. We demonstrated how pangenomics can be applied to design broadly-applicable outsourced markers, and integrate a range of best practices, including local stakeholder input, strong inference, and digital decision support software, to facilitate breeding trait-enhanced varieties for African smallholder farmers. In this case, we used backcross introgression as proof-of-concept for nascent programs, but the most effective long term approach would be a pangenomic forward breeding in locally developed elite germplasm^10,14^. Pangenomic forward breeding would combine: (1) foreground selection of trait-predictive markers for desired traits, (2) background selection of trait-predictive markers for must-have acquired traits, and (3) forward polygenic selection for yield using genomic prediction. This approach would require updated low- and mid-density genotyping tools, to distill pangenomic discoveries into low-cost services, and new decision support software, to optimize selection across complex marker-causative variant relationships, allelic heterogeneity, and genetic heterogeneity. Pangenomic forward breeding networks that incorporate local stakeholders knowledge^7,47^ have the potential to deliver locally- and regionally-adapted varieties of staple crops that meet the demand of smallholder farmers and other stakeholders in developing regions.

## Methods

### Plant materials

Two seed sources of the resistant genotypes SRN39 were used. The first source PI656027 was obtained from the United States National Plant Germplasm System (NPGS). The second source SRN39 was a product of one selfing of PI656027 (from NPGS) in our program. Biparental families were developed with SRN39 as the donor parent while susceptible parents used were: Mota Maradi (PI656050), MR732 (PI656051), and SEPON82 (PI656024). All four parents are released varieties in Niger that are included in the sorghum association panel^15,50^. We confirmed that the recurrent and donor parents carried the expected alleles (*LGS1* susceptible and *lgs1-1* resistant, respectively) using the published PCR-based assay^29^. Briefly, genomic DNA of BTx623, Mota Maradi, El Mota, and SRN39 was extracted following the CTAB method and quantified using a NanoDrop spectrophotometer (Thermo Scientific). After normalization to 10 ng/μl, 2 μl of each sample was used for PCR. The first primer (G2133) has a predicted amplicon size of 351 bp on all genotypes assayed, while the second primer (PDstrigalgs5b) has a predicted amplicon size of 628 bp for only susceptible genotypes without *LGS1* deletion.

### Population genomic analyses for *lgs1* marker design

GBS reads were aligned to the BTx623 sorghum reference genome version 3.1^56^ using BWA-MEM^57^ and the TASSEL 5 GBS pipeline^58^ was used to call SNPs. We targeted a ∼0.5 Mb upstream flanking region (UFR; 69,400,000–69,977,147|v3.1; 577 kb) and a 100 kb downstream flanking region (DFR; 70,011,172–70,111,172|v3.1) on either side of the *lgs1-1* deletion interval to identify candidate SNPs for conversion to KASP markers. The smaller DFR interval was chosen to account for the very high subtelomeric recombination rate on the right arm of Chr 05 in sorghum^15^ which could increase the risk of uncoupling of causative and marker allele during breeding. A frequency of less than 80% missing data within the region was considered to account for polymorphism, not related to the deletion, present in the flanking regions. The extent of LD between the inferred deletion and flanking SNPs was quantified using *R*^2^ and *D*′ in TASSEL 5 software^59^. Allele frequencies were calculated using version 0.1.13 VCFtools^60^. SNPs in LD with the deletion were selected to develop KASP markers. Only SNPs that distinguish between the resistance line, SRN39, and the susceptible lines (e.g. BTx623) were selected to design KASP primers and SNP verification. A total of 10 SNPs within the UFR and DFR were used to design primers with 50 bp flanking sequences around each SNP obtained from https://phytozome.jgi.doe.gov for *Sorghum bicolor* v3.1, based on the physical position (Fig. 2a, Supplementary Table 5) ranging between 41 to 534 kb upstream and downstream of the Sobic.005G213600 gene model. SNP verification of the 10 SNPs was conducted at Intertek-AgriTech (Alnarp, Sweden) to test that the putative SNPs selected are of good quality and differentiate between BTx623 and SRN39 (Extended Data Fig. 4). After the SNP verification, selected markers (based on the SNP quality assessment) were tested in inbred lines and progenies derived from resistant and susceptible lines to determine how the markers classify individuals based on a priori predictions on the inbreds and parents used for biparental families. We mapped georeferenced African germplasm^23–25,61,62^ for the deletion region on chromosome 5 (*N*_SNPs_ = 21,926). A frequency of less than 80% missing data within the region was considered to account for allelic series *LGS1* referring as absence or presence of the deletion. Within the *lgs1-1* interval, the number of SNPs is 21 in BTx623|v3.1, and SRN39 has all 21 missing SNPs. An expanded GBS dataset^62^ was used to generate the allele map (Fig. 2c), containing 30 SNPs within the same deletion interval due to higher coverage.

### KASP genotyping assay for *lgs1*

Leaf tissues were collected from 14 day old plants grown in the greenhouse. A circular ¼ inch (∼ 6 mm) hole punch was used for tissue collection. Two leaf punches of each sample were placed into the respective well of a 96-well plate. Each accession has at least three individuals referred to as biological replicates. Samples from individual plants of the same seed lot were considered biological replicates, while samples from the same plant were considered technical replicates. “Artificial heterozygote” samples were generated by mixing leaf tissue from one leaf punch of the known resistant genotype with one leaf punch of the known susceptible genotype and used as positive controls for heterozygous genotype calling. Other inbred lines and progenies (F_2:3_) were replicated either four or two times in the plate. Each plate contained three blank wells; one randomly assigned well and the last two wells (H11 and H12) were left empty as negative controls. To dry the leaf samples, the plate was placed in a plastic bag containing silica beads (“Dry & Dry” Premium orange indicating silica gel beads, Dry & Dry. Silicagel Factory, Brea CA) overnight at room temperature. Dried leaf tissues were sent for DNA extraction and KASP genotyping at Intertek AgriTech (Alnarp, Sweden). Samples were randomized on each genotyping plate. The biparental families of Mota Maradi × SRN39, SEPON82 × SRN39, MR723 × SRN39 (F_2_ progeny) were grown in the greenhouse and tissue was collected 14 days after planting. F_3_ progenies of Mota Maradi × SRN39 were grown in the field from June to October 2019 in Manhattan, Kansas. For each F_2:3_ line, 50 seeds were sown within the plot. For the remaining F_2:3_ lines, a single plant was genotyped. Using the markers, backcross lines were obtained using Mota Maradi, MR732, and SEPON82 as recurrent parents. Near isogenic lines (both positive and negative at snpSB00246) were selected. All experiments were done on ILs derived from BC_3_F_7_ and F_2_:BC_1_F_7_. KASP genotyping was also conducted from individual plants in the field in Niger. Note, the lgs1 MABC material from INRAN was not included in the network-wide genotyping.

### *Striga* germination stimulant assay

The germination assay was conducted in the quarantine lab in the Department of Biology at Pennsylvania State University following a published protocol for *Striga hermonthica* germination assay^36^. Briefly, *Striga hermonthica* seeds, collected in Siby, Mali (12°23′ N; 8°20′ W), were used for the germination assay. *Striga* seeds were sterilized with 0.5% sodium hypochlorite for 30 seconds and rinsed three times with sterile distilled water. Approximately 75 seeds (counted under a microscope) were transferred into a 12-well plate with 500 µl of deionized water (DI) in each well. Seeds were preconditioned in the dark for 10 days at 30℃ before root exudate application. Progenies (BC_3_F_3_ and F_2_:BC_1_:F_3_) derived from SRN39 and Mota Maradi, MR732, and SEPON82 (*n* = 19) were used for the germination assay. Sorghum seeds were grown in sand and calcined clay mix (1:1 sand/calcined clay) for four weeks in the greenhouse. For each line, six biological replicates were grown. Plants were watered daily with fertigate (top-watering) with ¼ strength Miracle-Gro® Water Soluble All Purpose Plant Food (The Scotts Company, LLC. Marysville, OH). Four weeks after planting, each plant (root and shoot) was removed from the pot. Roots were washed with DI water to remove remaining potting debris. Plants were placed in a flask containing DI water using a 1:5 ratio of root:DI water, sealed with parafilm, and kept under darkness at room temperature for 48 hours. The solution was centrifuged at 4°C at 9,000 rpm for 10–15 minutes, until supernatant was clear, to remove residual root debris. Supernatant was placed on ice until the germination test. Root exudates (1.5 ml) were applied to preconditioned *Striga* seed. Three technical replicate assays were used for each biological replicate. The synthetic strigolactone GR24 (CAS Number 76974-79-3; ChemPep Inc, Wellington, FL) at 0.1 ppm and DI water were used as positive and negative controls, respectively. After 72 hours, germinated *Striga* seeds were counted under a stereomicroscope (Manufacturer: Amscope, specifications: 3.5–180X Manufacturing 144-LED Zoom Stereo Microscope with 10MP Digital Camera).

### Phenotyping for the *Striga* resistance program

#### Pot experiments

Pot experiments on *Striga* resistance of ILs were conducted in the research center at Niamey, Niger during the rainy seasons (June–October) in 2022, 2023, and 2024 (Extended Data Table 2). *Striga hermonthica* seeds used for the experiments were collected from *Striga*-infestation hotspots in smallholder farmer sorghum fields of Niger (Bazaga; 13.8° N, 5.1° E, ∼320 km east of Niamey). Soil for the pot experiments was collected near the station in Niamey. Two *Striga* seeds inoculations were performed. The first inoculation of *Striga* seed was conducted at sowing sorghum seeds while the second inoculation was conducted after pre-condition of *Striga* seeds for 11 days; at this time sorghum plants reached 2 weeks old. The number of emerged *Striga* were counted weekly from the apparition of the first striga plant in each pot in the trial until physiological maturity or death of the sorghum plant. Sorghum biomass (above- and below-ground) and *Striga* biomass (above-ground) were collected for each pot. *Striga* plant biomass from individual pots were harvested, dried at room temperature, and weighed. Sorghum ground biomass was harvested and dried at room temperature along with carefully washed sorghum roots from individual pots.

#### Field experiments

The field experiments (Extended Data Table 2) were conducted in Niger under artificial infestation, natural infestation, and uninfested control fields (*Striga*-free fields) in six locations on (BC_3_F_7_ and F_2_:BC_1_F_7_ progenies of MR732 and Mota Maradi, respectively). Planting was done during the rainy season in 2023 and 2024 (June–October). In artificial infestation (on-station, sick plot), sorghum seeds were sown with *Striga* seeds (∼1g/hole) collected from 10 Konni and Bazaga villages with no fertilizer amendment during the trial. There were no additional inputs in the natural infestation (farmer’s field). For all the locations, phenotypic data recorded are days to flowering, plant height, and yield components. For *Striga* related traits, the number of striga plants at 45, 60, and 90 days after sowing were recorded. Field trials in Togo were conducted under natural *Striga* infestation in Boufale (Kara Region) in 2023 and 2024.

### Mid-density genotyping assay

Development of mid-density amplicon-based genotyping assay (DArTag; Intertek Agritech, Sweden) for sorghum was initiated by collating sequencing data from 1,891 breeding lines collected from diverse programs across Asia, Africa, and international partners^63^. DNA was extracted from young seedling leaf tissue using a modified CTAB method and genotyped through Genotyping-by-Sequencing (GBS) with ApeKI enzyme^15,34^. SNPs were called using a hybrid pipeline combining GATK and BCFtools^64^, aligned to the *Sorghum bicolor* v3.1 reference genome^56^. After applying quality filters, mean depth ≥1, base quality ≥30, approximately 383,000 high-quality SNPs were retained. To construct the SNP panel, SNPs with >70% missing data were excluded, retaining only biallelic markers with MAF ≥5% and heterozygosity ≤5%. SNPs with high polymorphism information content (PIC > 0.25) were prioritized, and closely linked markers were removed. Marker specificity was validated via genomic alignment, and LD analysis was performed to ensure representative of markers across LD blocks. Functional annotation using SnpEff helped prioritize genic and near-genic SNPs. The final panel integrated criteria such as informativeness and trait relevance. Later we manually included curated SNPs available in literature linked to shoot fly, stay-green, Striga, aphid resistance, fertility restoration, and Al tolerance. Mid-density genotyping was used to test characterize the genome of the recurrent parent was recovered, Genotyping was conducted on introgression lines (BC_3_F_7_ and F_2_:BC_1_F_7_ progenies of MR732 and Mota Maradi, respectively) and as well as parent lines, as controls. Leaf tissues from each individual were collected in Niger with four replicates per genotype. Visualization and calculation of parent genome recovery percentages using the panGenomeBreedr package^35^.

### Marker development from resequencing

To ensure data integrity, we excluded 468 libraries flagged as duplicates by kinship analysis and high missing-data rates, resulting in a curated dataset of 1,676 libraries for downstream analyses. SNPs flanking the *LGS1* gene (600 kb upstream and 200 kb downstream) were extracted from the sorghum resequencing panel (*N* = 1676) using the Samtools HTSlib program^64,65^. To tag the *lgs1-1* deletion (34 kb) for LD analysis, an artificial SNP (S5_75078000|v5.1 A/T 99% reference vs. 1% favored allele) was introduced into the 1676 pangenome data set, with the T allele in lines with the *lgs1-1* allele (Framida, SRN39, 54.K.94.WitchweedRes, and IS10234). LD between S5_75078000|v5.1 and other SNPs was conducted using the TASSEL package^59^. Two SNPs (S5_75081793|v5.1, C/T, 1% MAF; S5_75132263|v5.1, G/A, 2% MAF) in LD with the introduced SNP were selected for KASP marker development at Intertek. Favored versus alternate alleles were differentiated through the competitive binding of two allele-specific forward primers. KASP marker optimization was conducted on leaf samples submitted from the Nigerien program. Data visualization of LD analysis and KASP genotyping results was conducted using the R plot function.

### Pangenome reference analysis

#### Pangenome graph

To identify structural variants across reference haplotypes (*N* = 33) around the *LGS1* region and visualize them in a tubemap, we built a pangenome graph using Minigraph-Cactus^66^ (v2.9.3) with default settings. Construction relied on the full v5.1 BTx623 Chr05 sequence^49^ as the primary reference and sequences for the remaining 32 haplotypes were subset to include the *LGS1* region plus 100 kb flanking sequence up- and downstream. For the *LGS1* region, we further processed this graph with vg toolkit^67^ and vcfwave (vcflib v1.0.10)^68^ to reduce allelic complexity and retain only variants ≥5 kb in length. Note, BTx623 is within the Macia haplotype group but distinguished in tubemap for illustrative purposes.

#### kmer genotyping

The kmer genotyping methodology^49^ extracts kmers (80mers in this instance) that are single copy within individual references and variable among members of the pangenome (*N*-1 references; *N* = 32). The pipeline proceeds as follows: (1) Each reference genome is individually converted into kmers, where each kmer and its frequency in the assembly is stored in a hash table. Any non-single copy kmers are flagged to be ignored during downstream analyses. (2) Individual reference hashes are combined in a single hash (main_Hash), updating the flags such that only single copy kmers among all considered references will be used downstream. To isolate kmers that were diagnostic of the *LGS1* haplotype structure, sequences were subsetted from the pangenome graph. Steps 1 and 2 above are repeated the subsetted sequences, generating a combined subsetted hash that contains local single-copy sequences from the *LGS1* region across the pan-genome. (3) To ensure the local single copy markers do not occur elsewhere in the genome (which would prevent their use as a genotyping marker), the combined *LGS1* subsetted hash is intersected with the main genome hash, updating flags such that any non-single copy kmers will be ignored when genotyping (genotyping_hash). (4) Illumina libraries (*N* = 2,145) are genotyped by searching for all kmers in the genotyping_hash and counting their frequencies. A priori clusters were generated based on the haplotypes at *LGS1* locus within the pangenome tubemap. Diagnostic kmers were sorted into non-overlapping bins from reference Illumina library kmer frequencies (Macia:21,545; Mota Maradi:10,966, IS12646:545, Ajabsido:10,414, SRN39:326, RTx430:80), then kmer matches within each resequencing Illumina library and reports a percentage match from each bin, regardless of which reference they were derived from (e.g. LibraryA|1400|2:93:1:3:1:0; where Library A matched 1400 total diagnostic kmers across six haplotype bins). Illumina library haplotypes were then assigned based on majority match (e.g. Haplotype 2 - 93% of matches) to haplotype bins.

#### LD analysis

For LD analysis of kmer calls, libraries were assigned a haplotype for *lgs1-1* dependent on the proportion of kmer hits to SRN39 hashes. Libraries whose largest number of unique kmer hits were from the SRN39 haplotype were assigned a “1” and all other libraries were assigned a “0”. This kmer-inferred *lgs1-1* call was used to calculate LD (*R*^2^ and *D*’) with all SNP and InDel variants flanking LGS1 (Chr05, 74850000–75350000) using PLINK1.9.

### Pangenome-enabled breeding for staygreen drought tolerance

#### Marker development and testing

Developing improved versions of farmers’ preferred varieties with increased drought tolerance, while maintaining acquired traits for acceptability, was a common goal of the ISRA-Senegal and INERA-Burkina Faso sorghum breeding programs. To achieve this goal within a five year timeframe, we took advantage of discoveries made by the international scientific community^28,69,70^ to identify *Stg1*, *Stg3A*, *Stg3B*, and *Stg5* markers for use in oligogenic MAS in ISRA and INERA breeding programs (Extended Data Fig. 3a). Markers at *Stg2* and *Stg4* were not included in the MABC where BTx642 (B35) was used as donor because their favorable alleles are harbored by local cultivars instead. We used the 1,676 whole genome resequencing panel^49^ and 1,362 existing KASP assays (LGC Biosearch Technologies, Middlesex, UK) to identify polymorphic markers at *Stg3A* and *Stg3B* genomic regions suitable to use and scalable across other programs, specifically INERA, ITRA, and INRAN. After validation, two markers at *Stg3A* (S2_56954578|v3.1, S2_70591587|v3.1) and two markers at *Stg3B* (S2_69739036|v3.1, S2_71419274|v3.1) were retained for use in marker-assisted backcrossing activities in the selected ISRA and INERA farmers’ preferred varieties. For *Stg1*, the KASP marker S3_66366589|v3.1, which is carried by BTx642, was used in ISRA and INERA population development. In addition, with the discovery of another stay-green *Stg5* locus which co-localizes to the dhurrin biosynthetic gene cluster and confirmed to contribute post-flowering drought tolerance^69^, additional KASP markers were developed.

#### Marker assisted selection

Following validation two flanking markers around *Stg5* (S1_01147523|v3.1, S1_01287238|v3.1) were used for MABC selection in the ISRA pilot program, then in INERA breeding programs (Extended Data Fig. 3b). Backcross breeding at ISRA consisted of one backcross for Nganda × BTx642 (advanced to the BC_1_F_8_) and three backcrosses for both Darou × BTx642 and Golobe × BTx642 (advanced to the BC_3_F_8_). After each backcross generation, progeny were genotyped and those carrying at least one stay-green QTL (*Stg1*, *Stg3A*, *Stg3B*, or *Stg5*) were retained. At INERA, the elite variety IRAT204 was crossed with BTx642. Three hundred F_2_ progenies were planted in the field at Kamboinse Research Station, and leaf samples were collected and sent for KASP marker genotyping at Intertek Lab, Sweden. Superior lines carrying staygreen alleles were marked and self-pollinated to produce F_3_ recombinant inbred lines (RILs) before being advanced for several generations to obtain a total of 168 F_5_ progenies (Extended Data Fig. 3g).

### Phenotyping for the drought tolerance program

#### Field trials

Two off-seasons managed drought stress experiments were performed at CNRA research station in Bambey (14°42′N; 16°28′W), Senegal in 2022 (Nov 2022-March 2023) and 2023 (Feb 2023-June 2023). A total of 30 NILs were used in the 2022 field experiment and 234 NILs in the 2023 field experiment. The 2023 managed drought stress field experiment was conducted during the hot off-season (February-June) at the CNRA research station in Bambey, Senegal (14°42′N, 16°28′W) (Extended Data Fig. 3d). The experiment included 234 BC_1_F_5_ and BC_3_F_5_ lines from ISRA, parental checks (Golobe, Nganda, Darou, BTx642) as well as reference checks (non-staygreen Tx7000 and drought tolerant CE145-66) for a total of 240 genotypes. A 20 × 12 triple alpha lattice design was used. Each plot consisted of two rows of 2 m in length separated by 0.7 m with 0.2 m between plants within a row. Four to six seeds were sown per hill, and seedlings were thinned to one plant per hill at approximately 20 days after sowing (DAS), achieving a planting density of ∼65,000 plants/ha. Two irrigation regimes were applied, well-watered (WW) from sowing to maturity and post-flowering water-stressed (WS), with irrigation withheld in the WS treatment when 75% of the entries (genotypes) had reached flowering until physiological maturity. A similar managed drought stress field experiment at CNRA-Bambey in 2022 is pictured in Extended Data Fig. 3f. At INERA-Kamboinse research station, Burkina Faso (12°27′N, 1°32′W), rain-fed field screening was conducted for the IRAT204 × BTx642 F_6_ progenies at the end of the growing in 2022 (Extended Data Fig. 3g). A total of 19 lines were selected based on agronomic performance, earliness, and grain hardness and vitreosity. The 19 lines were KASP genotyped, among which nine carry one or more *Stg* alleles. A final set of four lines was retained for field evaluation in three different stations and in the farmers’ fields surrounding each research station.

#### Pot trials

A managed drought stress pot experiment was carried out in the screenhouse at CERAAS in Thies, Senegal (14°45′N, 16°53′W) during 2024 hot off-season (March – June) (Extended Data Fig. 3c,e). A total of 16 genotypes (12 ISRA introgression lines and the four parental checks) were used, with three biological replications (pots) per line and three replications for each treatment (WW pots and WS pots) for a total of 288 plants. Water deficit was applied at 50% flowering of plants for a given genotype for a period of 10 days. SPAD chlorophyll content was measured at different time points including physiological maturity.

## Data availability

Reference genome assembly and annotation files of all pangenome members are available at https://phytozome-next.jgi.doe.gov/. All raw sequence reads have been deposited in the NCBI SRA database under BioProject accessions listed in the pangenome reference paper^49^. Genotype and phenotype data is available at https://github.com/CropAdaptationLab/PangenomicBreedingNetwork.

## Description of Supplementary Information

Supplementary Information | Document with Supplementary Tables 1-6 and Supplementary Data 1-3.

Supplementary Table 1 | Market Segment Descriptions for West and Central Africa

Supplementary Table 2 | Market Segment Descriptions for East and South Africa

Supplementary Table 3 | Target Product Profile Descriptions for West and Central Africa

Supplementary Table 4 | Target Product Profile Descriptions for East and South Africa

Supplementary Table 5 | Summary of genotyping for 10 candidate SNPs for lgs1 KASP markers

Supplementary Table 6 | KASP genotype calls of snpSB00246 in INRAN germplasm

Supplementary Data 1 | GBS data of Sorghum Association Panel and West African Sorghum Association Panel on Chr05

Supplementary Data 2 | kmer genotyping results for resequencing panel

Supplementary Data 3 | Linkage disequilibrium of resequencing variants with kmer-inferred lgs1-1 deletion

## Code availability

Scripts and analysis methods are available at https://github.com/CropAdaptationLab/PangenomicBreedingNetwork and https://github.com/awkena/panGenomeBreedr.

## Acknowledgements

This paper is made possible by the support of the American People provided to the Feed the Future Innovation Lab for Collaborative Research on Sorghum and Millet (SMIL) through the United States Agency for International Development (USAID) under Cooperative Agreement No. AID-OAA-A-13-00047. The contents are the sole responsibility of the authors and do not necessarily reflect the views of USAID or the United States Government. We thank the USAID-SMIL management entity and external advisory board for guidance on developing the network. The work was funded by the Gates Foundation through “Mining useful alleles for climate change adaptation from CGIAR gene banks (INV-030574)” (G.P.M., J.R.L.), “Accelerated Varietal Improvement and Seed Delivery of Legumes and Dryland Cereals in Africa” (H.G), and “Green Evolution - Accelerating Dryland Cereals Improvement for Africa (INV-053669)” (G.P.M., D.H.R.). The *Striga* resistance phenotyping in Niger was supported by TWAS-UNESCO. Portions of this manuscript were adapted from F.M.’s Ph.D. dissertation at Kansas State University. We thank colleagues who contributed to aspects of the network which were not addressed in this paper due to space limitations. The work (proposals: 503014; Award DOIs: 10.46936/10.25585/60001093) conducted by the U.S. Department of Energy Joint Genome Institute (https://ror.org/04xm1d337), a DOE Office of Science User Facility, is supported by the Office of Science of the U.S. Department of Energy operated under Contract No. DE-AC02-05CH11231.

## Author contributions

G.P.M., D.F., F.M., J.M.F, and H.G conceived and organized the research and development initiative. F.M., J.M.F., C.D., E.A., N.O., S.R.M., and T.J.F. conducted the molecular breeding activities. F.M., J.Ma., C.M.M, M.I.T., O.A.A. O.S.D., A.M.I., O.A.M., A.H., R.A., P.K., and E.A. conducted the *Striga* resistance experiments. J.M.F., J.P.S., S.B., M.N., D.S., B.S., C.D., and N.O. conducted the drought tolerance experiments. F.M., J.M.F., S.D., A.R., H.G., P.O.-O., S.R.M., G.P.M., and T.J.F developed the genotyping assays. F.M., J.M.F., A.M.H., A.L.H., C.M.M., J.T.L., P.O.-O., S.R.M., C.J.V., and C.C.-B. conducted population genomic and pangenomic analyses. A.W.K, I.T.T, B.A., F.M., J.M.F., G.P.M., L.B., and C.C.-B. developed panGenomeBreedR. F.M., J.M.F., C.D., O.A.A., O.S.D., A.M.I., E.A., N.O., M.A.Y., A.D., R.K., R.K.K., C.P.K., T.L., L.M., J.Mu., E.T.M., G.N., H.N.A.M., K.O.-O., R.O.A., S.R., T.S.A., and L.Y. contributed germplasm for network-wide genotyping. M.K.V.C., B.N., M.N.T, A.A., J.M.F., F.M., T.J.F., S.R.M., C.J.V., and H.G. coordinated and analyzed network-wide genotyping. G.P.M., F.H. D.F., A.M.I., F.M., J.M.F., C.D., B.S., E.A., N.O., A.W.K., J.T.L., T.J.F., J.R.L., C.C.-B, D.H.R, and H.G. acquired funding and supervised the research. F.M. and G.P.M. wrote the manuscript. J.M.F, S.R.M., J.T.L., A.R., and C.J.V. contributed text, figures, and/or revisions. All authors edited and approved the manuscript.

---

**Extended Data Fig. 1.**
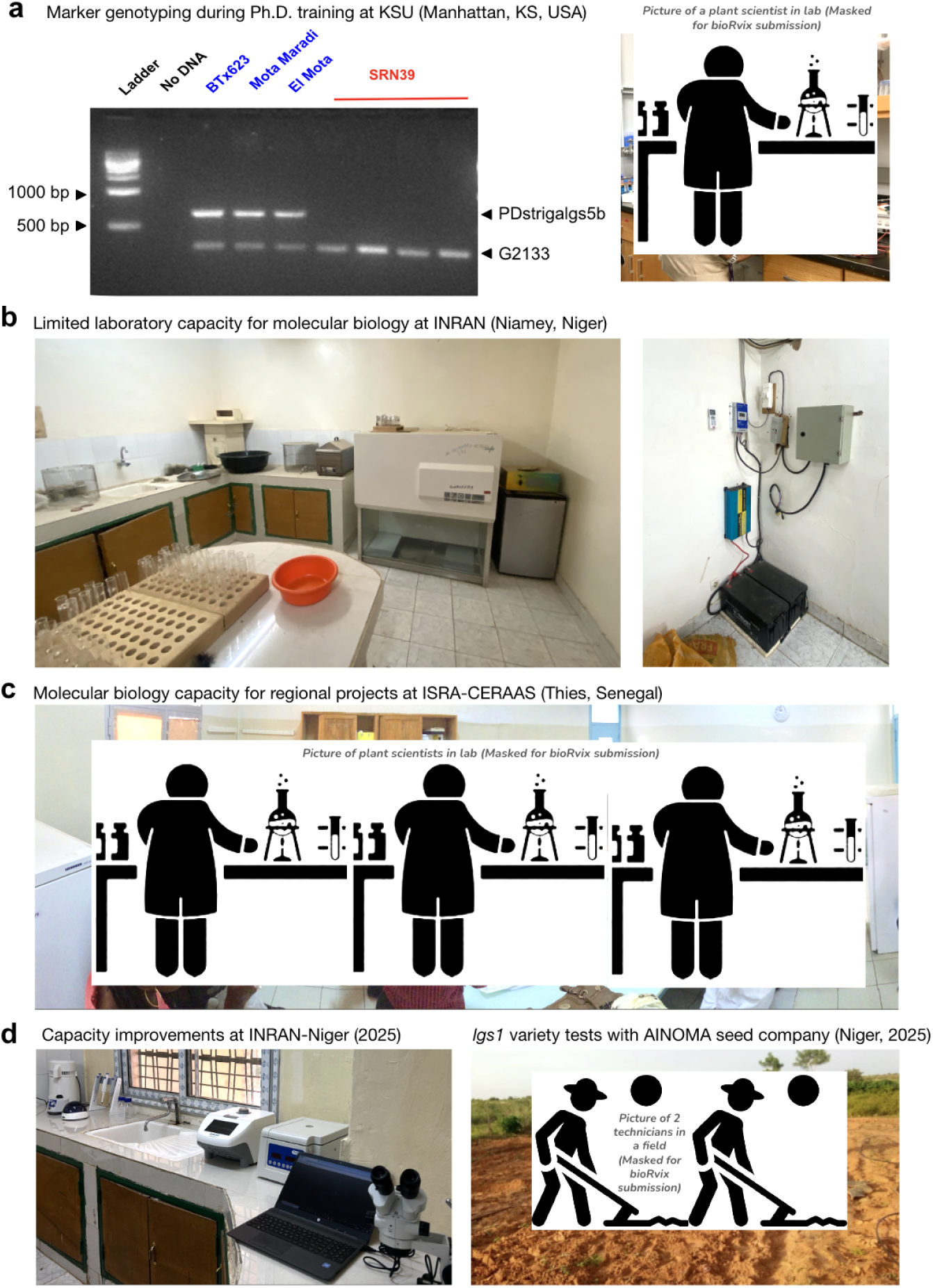
PCR markers are accurate in the INRAN breeding program, but are not tractable for MAS in Niger. **a**, INRAN attempted to introgress LGS resistance into locally-preferred varieties in the 2000s^22^, but was not completed due to various resource constraints. We relaunched the program, confirming the LGS1 status of INRAN breeding materials using two previously developed primers sets^29^ were adapted for a duplex reaction. G2133 (non-deletion target) represents the common primer that amplifies all samples while PDstrigalgs5b (deletion target) represents the LGS1 deletion primer that amplifies samples without deletion at *LGS1* gene (Sobic.005G213600). Ladder: 500 bp bands ladder; No DNA: negative control. BTx623, Mota Maradi, El Mota: *Striga* susceptible genotypes; SRN39: *Striga* resistant genotype. This experiment was done by F.M. during PhD training at Kansas State University (right panel; Photo: G.M.). **b**, INRAN-Niger is an example of an African NARI with limited capacity for molecular biological assays. At the launch of this initiative, the INRAN laboratory (left panel, Photo: G.M.) had some basic lab equipment (i.e. refrigerator, water bath, weigh scale, etc) but lacked standard molecular biology equipment (i.e. microcentrifuges, pipettors, spectrophotometer, PCR machine, gel electrophoresis system, ultra-low freezers) necessary for DNA extraction and PCR marker analysis and power outages were frequent at the facility and necessitated a battery backup system (right panel). **c**, ISRA-CERAAS in Senegal is an example of a NARI with high capacity with a dedicated molecular laboratory (Photo: G.M.) and other laboratories (for biochemistry, ecophysiology, nutritional analysis, bioinformatics, tissue culture, etc.). **d**, Recent investments (TWAS-UNESCO, FAO) have enhanced laboratory capacity at INRAN (e.g. microcentrifuge, pipettors, spectrophotometer, microscope, screenhouse; Photo: F.M.) so that INRAN scientists can contribute lab research activities within the pangenomic breeding network, in addition to critical field studies, such as those with local smallholders and/or **e**, local seed enterprises (AINOMA seed company trial, 2025; Photo: Aichatou Nasser).

**Extended Data Fig. 2.**
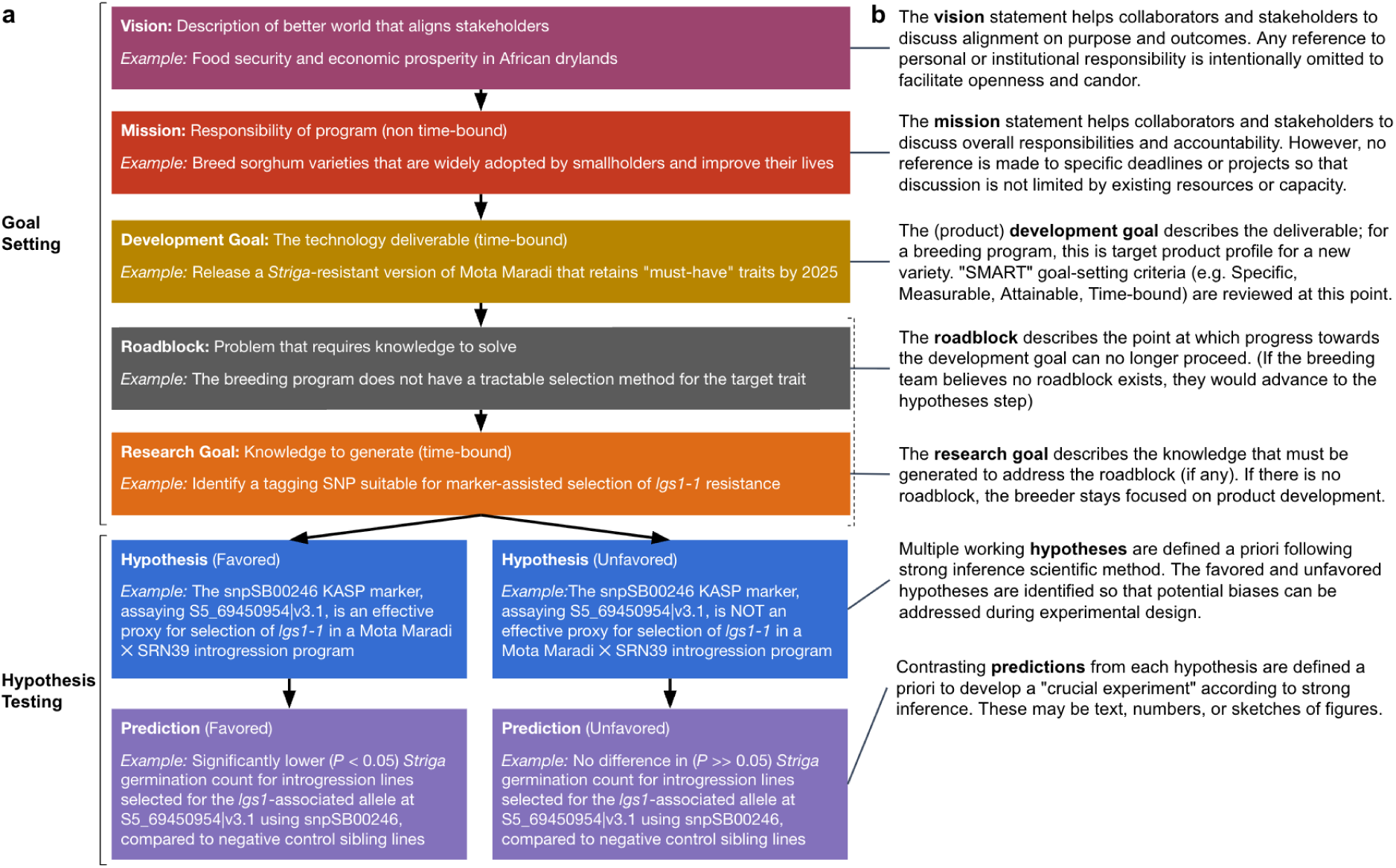
Framework for formalization of goal setting and hypothesis testing. **a**, An example of logic chains that formally link breeder goal-setting and strong inference scientific methods. **b**, Rationale for each step and description of how each step is completed. The goal-setting logic chain (upper five steps) adapts one provided by hfp consulting (Heidelberg, Germany), adding the distinction between “development goals” (for technology) and “research goals” (for knowledge), and adding the “roadblock” identification to help breeders remain focused on research activities that are essential to breeding goals. When the breeding team believes no roadblock exists, the roadblock and research goal may be omitted, such that development goals are linked directly to hypothesis testing. The development goal or research goal is linked directly to hypothesis testing. The hypothesis-testing logic chain (lower four steps, with branching structure) is adapted from the strong inference approach to the scientific method^32,71^, which emphasizes the use of multiple working hypotheses, the awareness of the tendency to favor some hypotheses, and the need to design “crucial” (decisive) experiments. Our approach extends strong inference by (1) explicitly naming the “favored” and “unfavored” hypotheses, so that the team can address potential biases during experimental design, and (2) facilitating the design of crucial experiments by explicitly defining predictions under each hypothesis. Templates and training materials are available at www.gohy.org.

**Extended Data Fig. 3.**
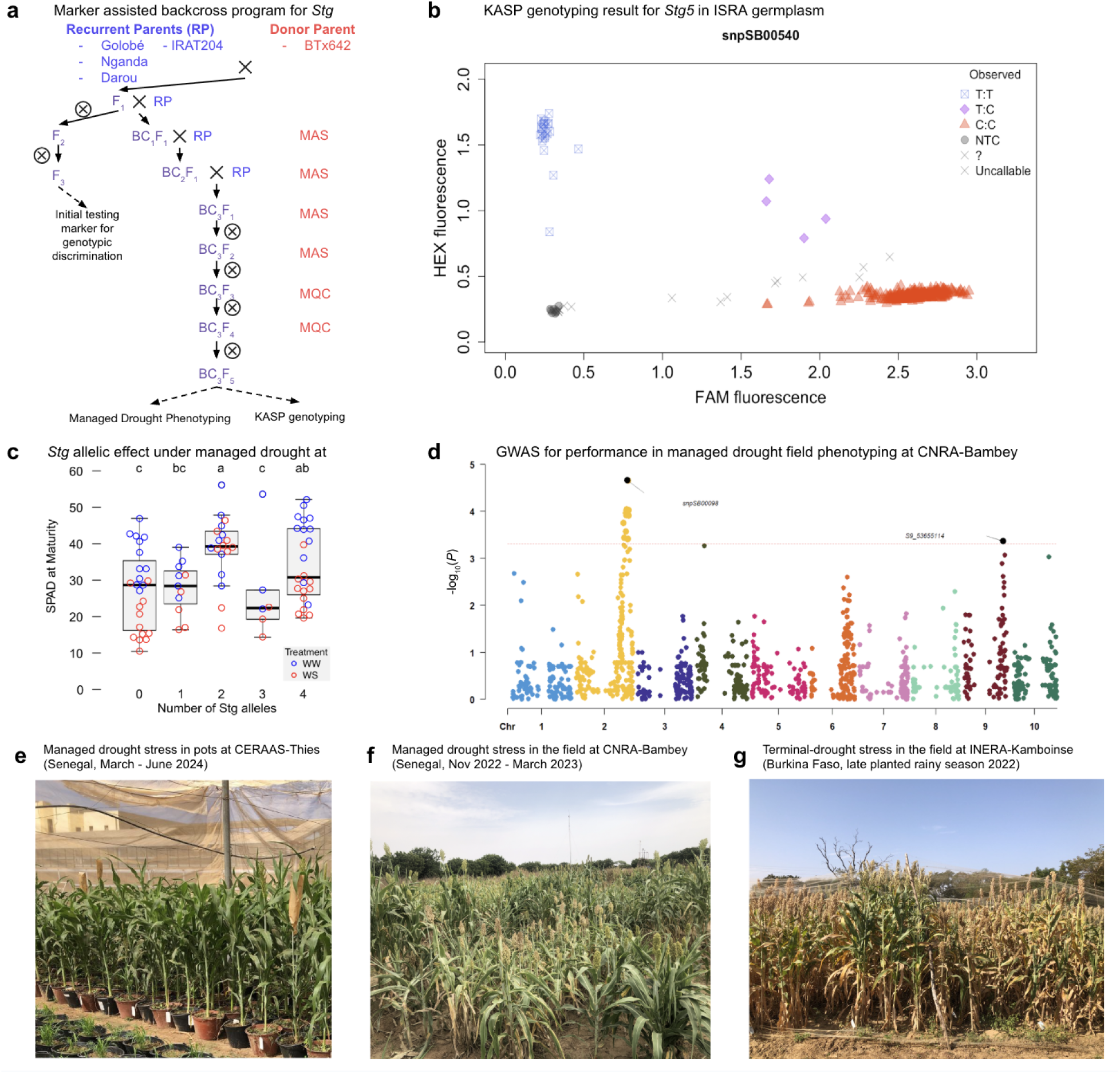
Pangenome-enabled breeding for staygreen drought tolerance at ISRA-Senegal and INERA-Burkina Faso. Marker assisted selection for staygreen alleles (*Stg1*, *Stg3A*, *Stg3B*, and *Stg5*) in locally-preferred varieties and managed drought phenotyping in Senegal and Burkina Faso. **a**, Steps of marker-assisted selection (MAS) and marker-based quality control (MQC) with *Stg* KASP markers. **b**, *Stg5* KASP marker (S1_01147523|v3.1, snpSB00540) scatterplot distinguishing segregating alleles in ISRA germplasm. **c**, Boxplot of drought response (SPAD) for BC_1_F_5_ and BC_3_F_5_ plants at physiological maturity conducted in a screenhouse at CERAAS-Thies in 2024. BC_1_F_5_ and BC_3_F_5_ lines were phenotype under well-watered (WW) and water-stressed pots (WS). The letters above indicate Tukey’s HSD. The number of Stg alleles, determined by KASP genotyping, indicates any combination of *Stg1*, *Stg3A*, *Stg3B*, and *Stg5* and demonstrated staygreen trait introgression to elite varieties. **d**, Genome-wide association study of drought stress index for grain weight identified loci contributing drought tolerance in the field at CNRA-Bambey during the 2023 off-season (February-June). Introgression lines from three backgrounds (Golobe, Nganda, Darou) were used, confirming the role of *Stg* and other loci in staygreen trait introgression. **e**, Post-flowering drought stress phenotyping of introgression lines in pots at CERAAS-Thies research station in 2024, data shown in panel c. **f**, Managed drought stress phenotyping representative of data in panel d at CNRA-Bambey during the 2022 late rainy season (Nov 2022-March 2023). BC_1_F_3_ and BC_3_F_3_ plants reached grain fill and maturity under post-flowering drought. **g**, Managed drought stress phenotyping at INERA-Kamboinse research station during the late growing season of 2022. Phenotyped RILs were enriched for Stg alleles by MAS.

**Extended Data Fig. 4.**
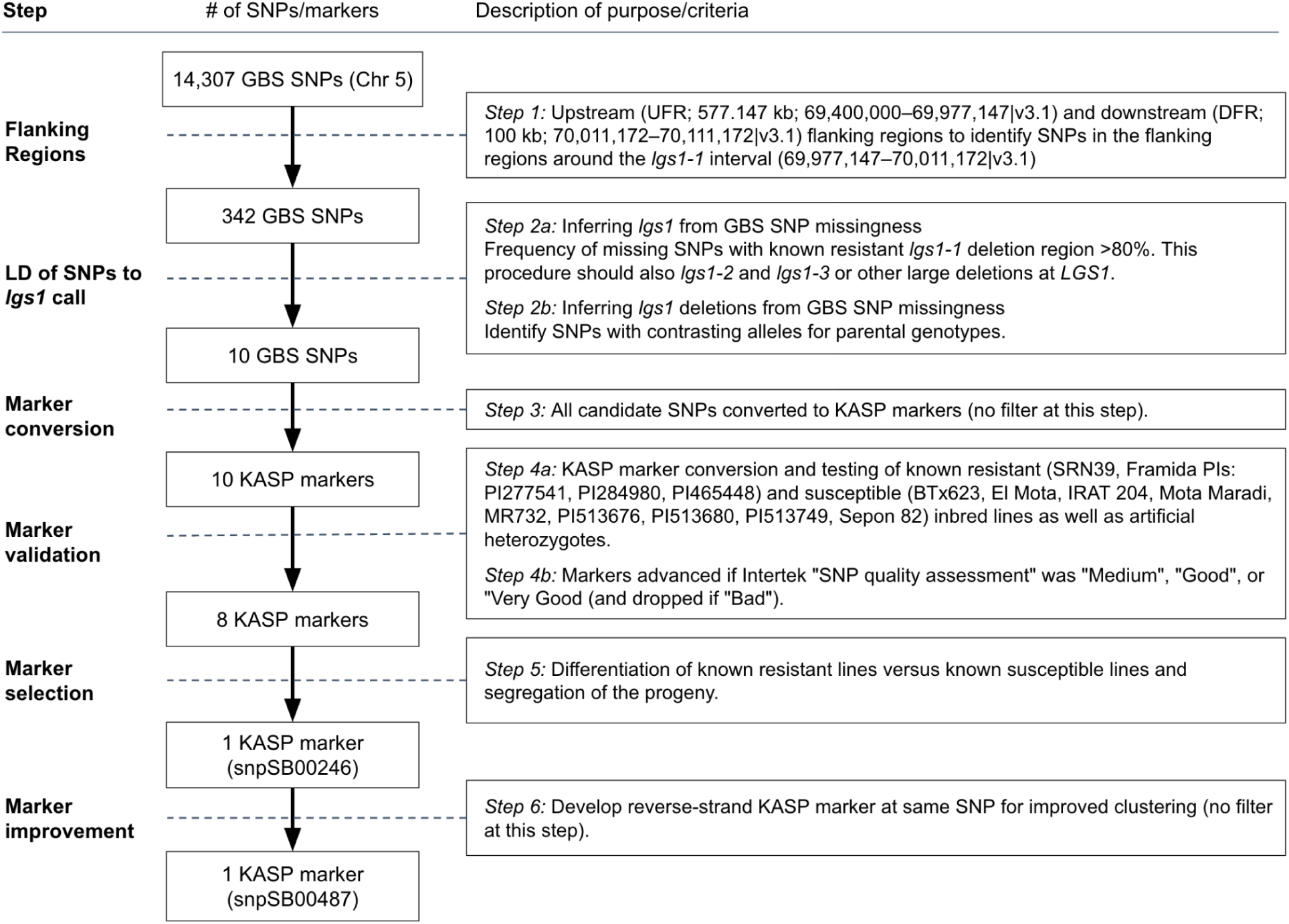
Workflow for development of outsourced KASP marker. Description of steps towards development of KASP markers based on linkage disequilibrium of SNPs with GBS-inferred *lgs1* deletion calls.

**Extended Data Fig. 5.**
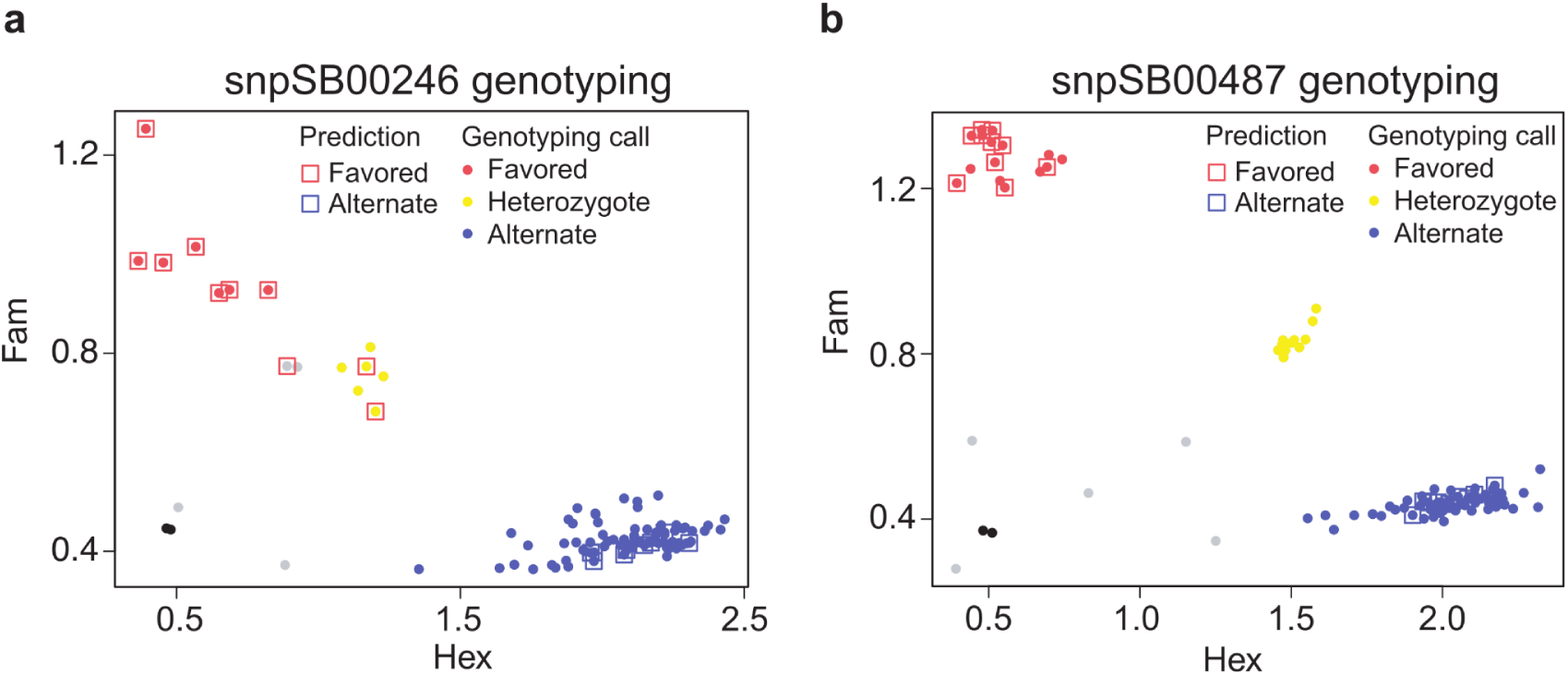
KASP marker generated on the reverse strand improved genotyping calling. **a**, KASP marker snpSB00246, generated on the forward strand sequence of SNP S5_74564291-v5.1, showed incorrect calling of homozygous favored allele as heterozygotes due to close clustering of heterozygous and the favored allele clusters in the ITRA-Togo sorghum breeding program in 2021. **b**, KASP marker snpSB00487, generated on the reverse strand sequence of SNP S5_74564291-v5.1, showed three distinct clusters. Genotyping calls of the favored and alternate alleles matched the allele prediction, indicating improved marker prediction in the ITRA-Togo program in 2021.

**Extended Data Fig. 6.**
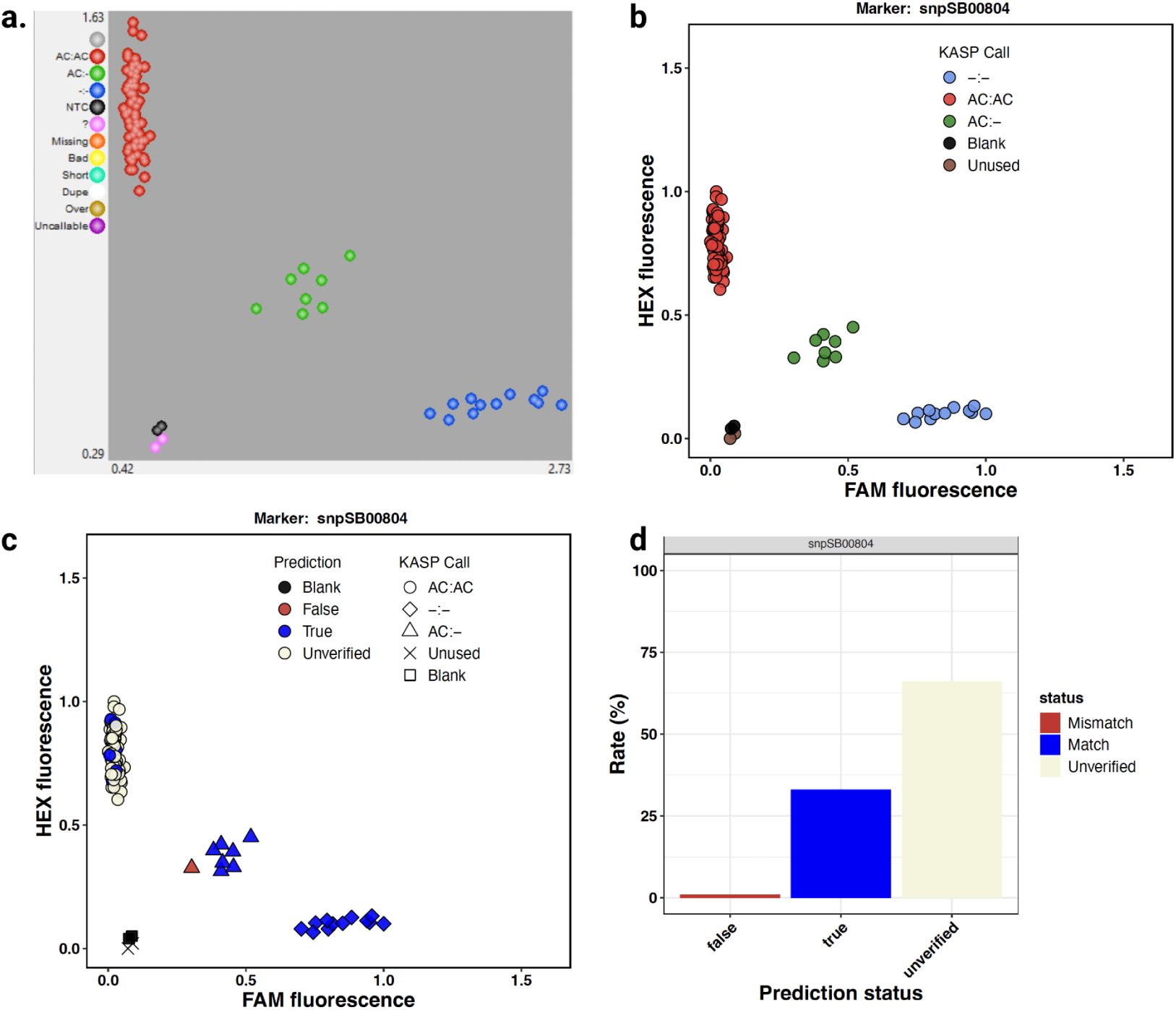
Incorporating strong inference principles into a decision support software for breeders (panGenomeBreedr). Comparison of allele/genotype discrimination cluster plots from **a,** standard software (SNPViewer) versus **b-d**, South-led decision support software (panGenomeBreedr) that incorporates strong inference. **a**, SNPViewer provides basic visualization with post hoc color-coding so erroneous genotype calls cannot be distinguished. **b**, By contrast panGenomeBreedr offers advanced functionality for accurate genotyping hypothesis testing by incorporating *a priori* predictions of known positive controls. **c**, Genotypes are represented using distinct symbols for inferred genotypes (homozygotes for each of two classes and heterozygotes). Notably, however, to facilitate hypothesis testing color overlays indicate the concordance between predicted and observed KASP genotype calls. Blue denotes concordant (accurate) genotypes, red indicates discordant calls (genotyping errors), and beige represents non-control samples for which verification is not possible. **d**, An additional barplot summarizes the genotype call accuracy across samples for the marker-plate combination to facilitate breeder decisions. Of the 94 samples, 32 were designated as positive controls; among these, 31 were accurately genotyped (blue) and 1 exhibited a genotyping error (red) (Panels c and d).

**Extended Data Fig. 7.**
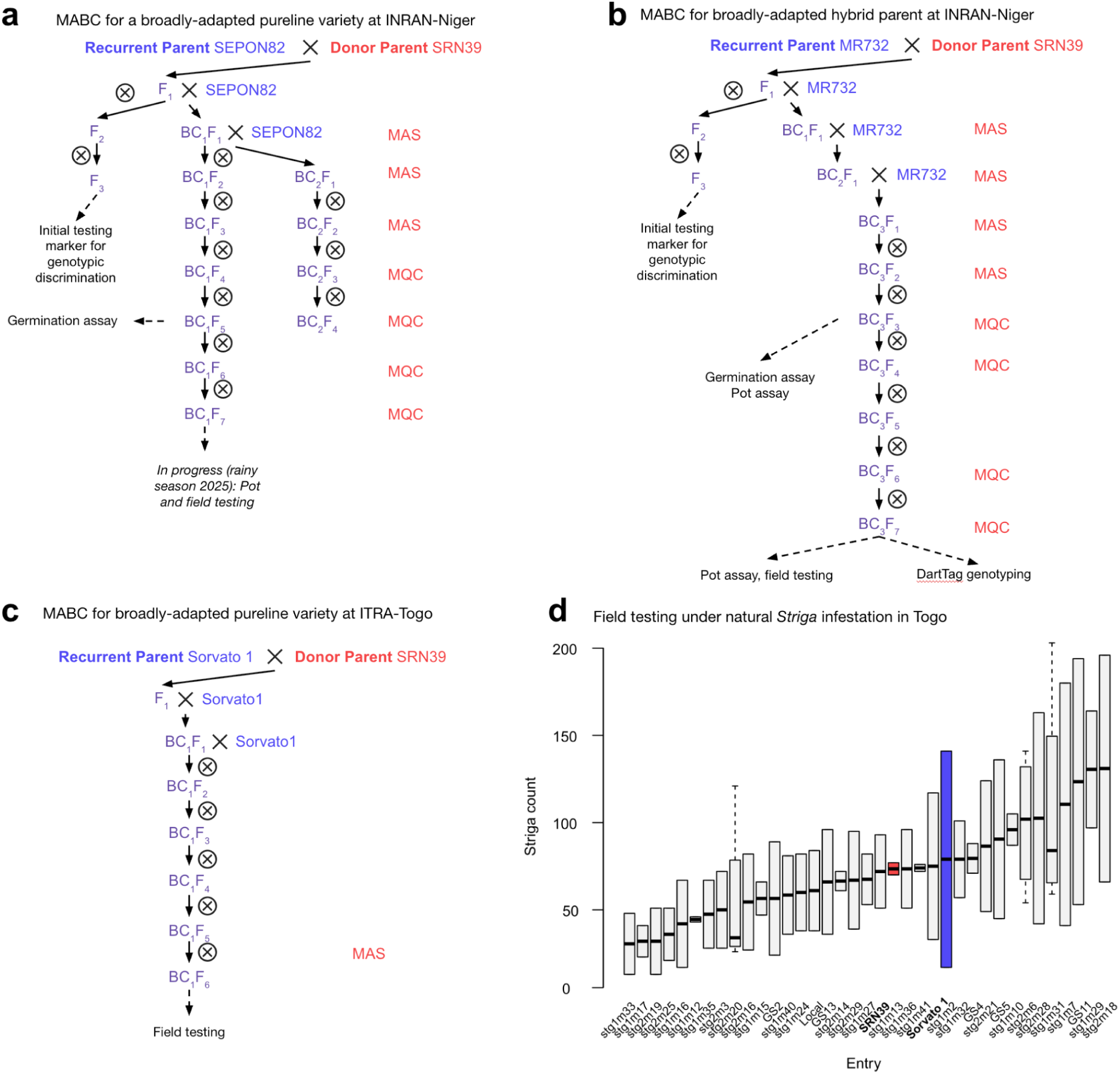
Marker-assisted introgression programs at INRAN-Niger and ITRA-Togo. Marker-assisted backcrossing of *lgs1-1* deletion allele and testing for **a**, SEPON82 at INRAN-Niger **b**, MR732 at INRAN-Niger and **c**, Sorvato1 at ITRA-Togo. Steps of marker assisted selection (MAS) and marker assisted quality control (MQC) are noted. **d**, Field phenotyping of *Striga* infestation counts of *lgs1* ILs and parent lines in naturally-infested *Striga* hotspot in Boufale, Togo (2023–2024). Note, this trial within the ITRA-Togo breeding program includes only *lgs1* ILs and not *LGS1* sibling ILs.

**Extended Data Fig. 8.**
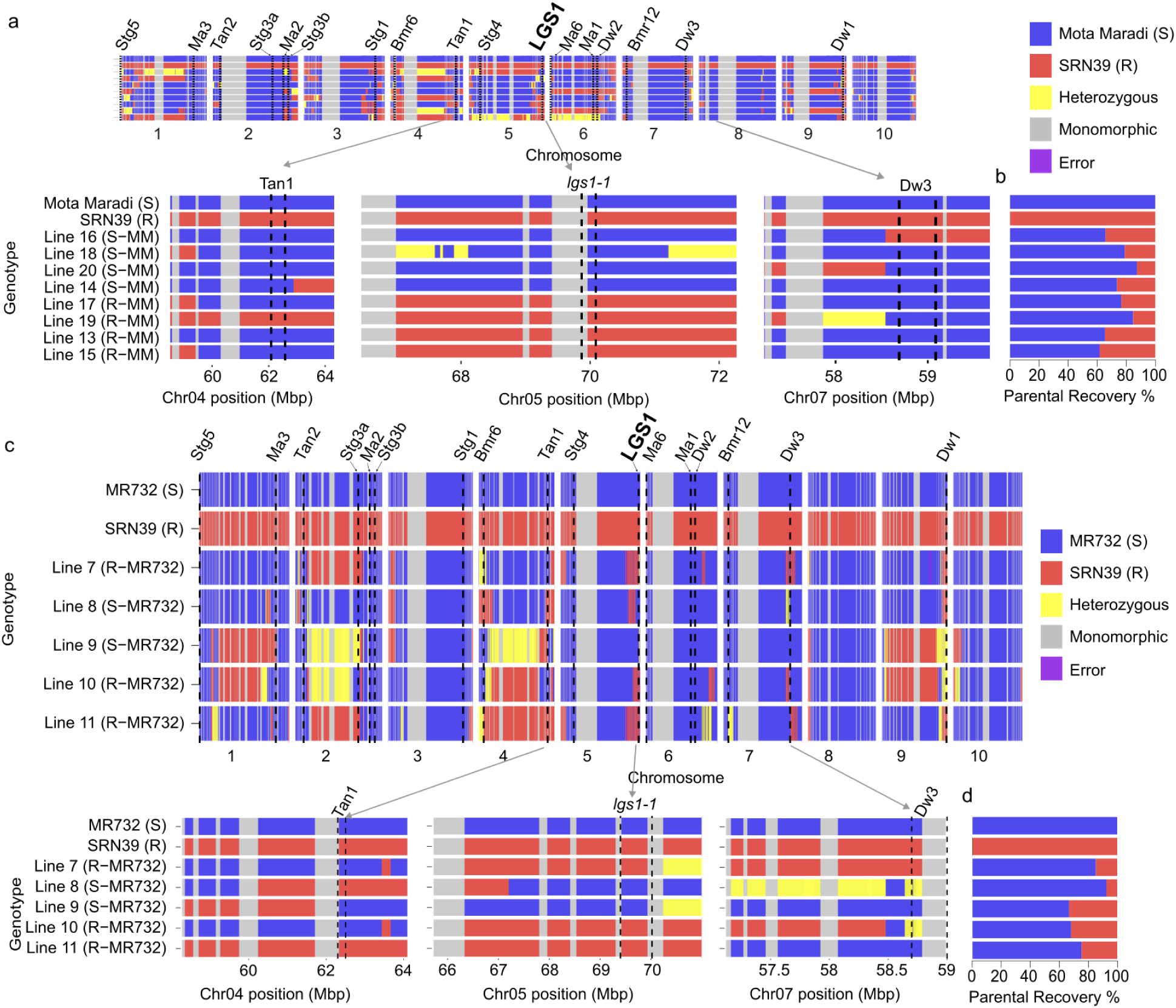
Genome-wide characterization of introgression lines guides pangenomic breeding strategy. Visualization of mid-density genotyping (DArTag) using panGenomeBreedr. **a**, Detailed view of selected acquired and desired trait loci in Mota Maradi. For instance, for the major height gene *Dwarf3* ^72,73^, Mota Maradi carries the wild-type allele (*Dw3*; tall) while the SRN39 carries the semi-dwarf allele (*dw3-ref*) that reduces forage yield. In this case, we were fortunate to recover the Mota Maradi haplotype in all the *lgs1-1* ILs, but a recently developed pangenomic-derived KASP marker^73^ could have easily ensured this outcome. A contrasting case is illustrated at *Tannin1*, the major gene conditioning grain tannins, one the most important local adaptation traits^74,61^. At *Tannin1*, one of the *lgs1-1* ILs had an unintended introgression of the SRN39 haplotype, which might suggest this IL would be unacceptable as a variety. However, we know from the pangenomic characterization that Mota Maradi and SRN39 are isogenic at *Tannin1* (*tan1b* carriers), so MAS at this locus would not only have been unneeded, but would have unnecessarily limited genetic gain from segregation on this chromosome. **b**, An average 72% recurrent parent genome was recovered in Mota Maradi *lgs1-1* ILs similar to the negative sibling ILs (77%, *P* = 0.56) and the expectation in the absence of selection (75%) (*t*-test *P* = 0.62). **c**, Detailed view of select acquired and desired trait loci in MR732. **d**, An average 76% recurrent parent genome was recovered in MR732 *lgs1-1* ILs similar to negative sibling ILs (80%, *P* = 0.79).

**Extended Data Fig. 9.**
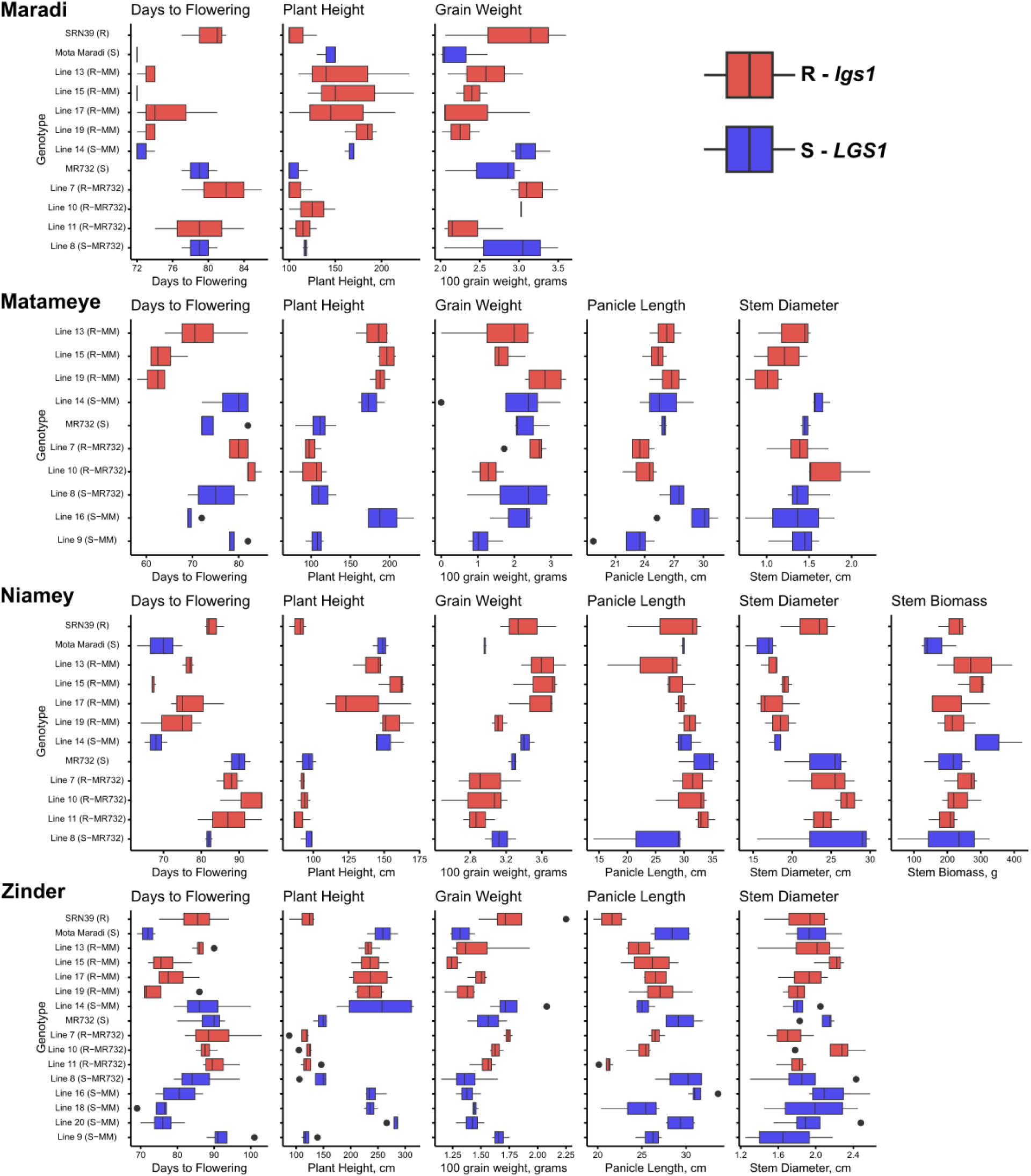
Field phenotyping trials on recovery of farmer-preferred traits. Box plots of flowering time, plant height, grain weight, panicle length, stem diameter and biomass evaluated on-station (Maradi, Niamey) or smallholder farmer fields (Matameye, Zinder) in Niger.

**Extended Data Fig. 10.**
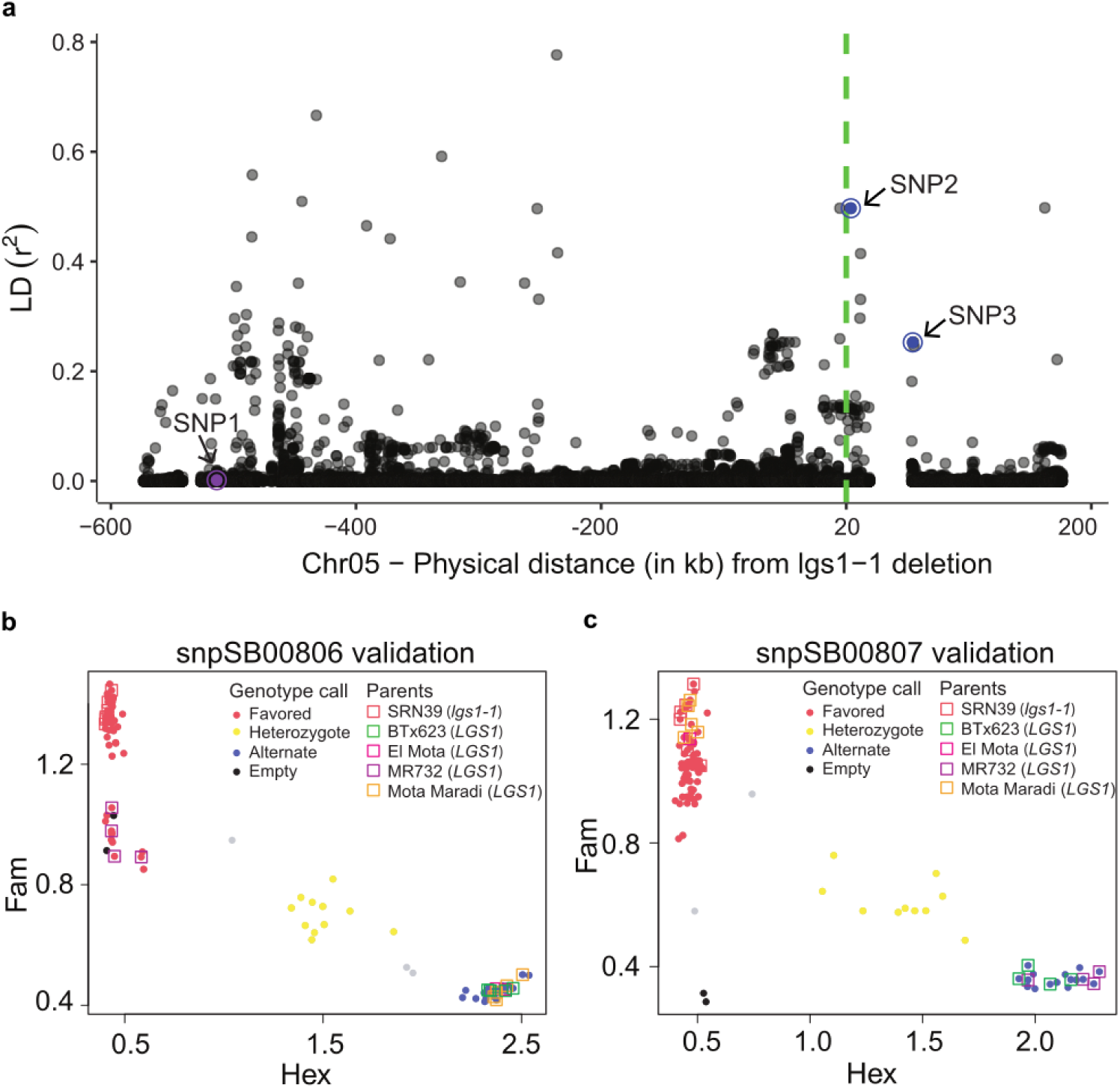
Pangenome-based development of markers suitable for introgression of *lgs1-1* into a broad range of elite African varieties. **a**, Linkage disequilibrium of variants proximal to the *lgs1-1* deletion inferred by manual inspection of deletion. SNP1 (snpSB00246/S5_69450954|v3.1), SNP2 (snpSB00806/S5_75081793|v5.1), SNP3 (snpSB00807/S5_75132263|v5.1). **b**, KASP genotyping of snpSB00806 in the Niger sorghum breeding program. Marker differentiated the favored (*lgs1-1* associated) vs. alternate allele accurately in BTx623, El Mota, and Mota Maradi, but failed in MR732 due to a large deletion surrounding S5_75081793|v5.1. **c**, KASP genotyping of snpSB00807 in the Niger sorghum breeding program. Marker differentiated the favored (*lgs1-1* associated) vs. alternate allele accurately in BTx623 and MR732 but failed in El Mota and Mota Maradi as the SNP at S5_75132263|v5.1 was conserved between SRN39, El Mota, and Mota Maradi.

**Extended Data Table 1:** Institutions participating in pangenomic breeding network.

| Institution<br>Abbreviation | Full name of institution<br>(Center at institution, if applicable) | Location | Type <sup>a</sup> | Role in network<br><sup>b</sup> | Projects <sup>c</sup> | Authors |
| --- | --- | --- | --- | --- | --- | --- |
| CSIR | Council for Scientific and Industrial Research<br>(Savanna Agricultural Research Institute) | Tamale, Ghana | NARI | VD | GE, AVISA | KOO |
| EIAR | Ethiopian Institute of Agricultural Research | Addis Ababa,<br>Ethiopia | NARI | VD | AVISA | TL |
| IER | Institut d'Economie Rurale (Centre Régional de<br>Recherche Agronomique) | Bamako, Republic of<br>Mali | NARI | VD | AVISA | AD |
| INERA | Institut de l'Environnement et de Recherches<br>Agricoles | Ouagadougou,<br>Burkina Faso | NARI | VD | SAWAGEN, AVISA | NO, CPK |
| INRAN | Institut National de la Recherche Agronomique du<br>Niger | Niamey, Niger | NARI | VD, NC, PR | SAWAGEN, AM,<br>GE, AVISA,<br>TWAS-UNESCO | FM, MIT, OAA,<br>OSD, AMI |
| ISRA | Institut Sénégalais de Recherches Agricoles (Centre<br>Nationale Recherches Agronomiques; Centre d'Étude<br>Régional pour l'Amélioration de l'Adaptation à la<br>Sécheresse) | Bambey and Thiés,<br>Sénégal | NARI | VD, TD, NC | SAWAGEN, AM,<br>GE, AVISA, ILCI | JMF, CD |
| ITRAD | Institut Tchadien de Recherche Agronomique pour le<br>Développement | N'Djamena, Chad | NARI | VD | AVISA | GN |
| ITRA | Institut Togolais de Recherche Agronomique | Lomé, Togo | NARI | VD | SAWAGEN, AVISA | EA |
| IAR | Institute for Agricultural Research | Samaru, Nigeria | NARI | VD | AVISA | ROA, MAY |
| IRAD | Institute of Agricultural Research for Development | Yaoundé, Cameroon | NARI | VD | AVISA | HNAM |
| IIAM | Instituto de Investigação Agrária de Moçambique | Maputo,<br>Mozambique | NARI | VD | AVISA | JM |
| KALRO | Kenya Agricultural and Livestock Research<br>Organisation | Nairobi, Kenya | NARI | VD | AVISA | RKK |
| TARI | Tanzania Agricultural Research Institute | Dodoma, Tanzania | NARI | VD | AVISA | ETM |
| ZARI | Zambia Agriculture Research Institute | Lusaka, Zambia | NARI | VD | AVISA | LM |
| ARC | Biotechnology and Biosafety Research<br>Center, Agricultural Research Corporation | Shambat, Sudan | NARI | VD | AVISA | TSA |
| NARO | National Agricultural Research Organization | Soroti, Uganda | NARI |  | AVISA | RK |
| KU | Kenyatta University | Nairobi, Kenya | AU | TD | AVISA | SR |
| KNUST | Kwame Nkrumah University of Science and<br>Technology | Kumasi, Ghana | AU | TD, PR, NC | GE, ILCI | AK |
| DARS | Department of Agricultural Research Services | Lilongwe, Malawi | AU | VD | AVISA | LY |
| UAS | Université André Salifou de Zinder | Zinder, Niger | AU | TD | TWAS-UNESCO | OAM, AH, RA |
| CIMMYT | International Maize and Wheat Improvement Center | Nairobi, Kenya;<br>Thies, Sénégal | NPRO | NC, TD, PR | AM, GE | JMF, MKVC, BN,<br>MNT, AA, HG |
| CIRAD | Centre de coopération internationale en recherche<br>agronomique pour le développement | Montpellier, France | NPRO | NC | SAWAGEN | DF |
| ICRISAT | International Crops Research Institute for the<br>Semi-Arid Tropics | Patancheru, India;<br>Sadoré, Niger | NPRO | NC, TD, PR | SAWAGEN | SD, FH |
| HA | HudsonAlpha Institute for Biotechnology | Huntsville, AL, USA | NPRO | PR | GE | CMM, AMH,<br>ALH, JTL |
| CSU | Colorado State University | Fort Collins, CO, USA | LGU | TD, PR, NC | SAWAGEN, AM,<br>GE, ILCI | CCC, CJV, AK,<br>GPM |
| KSU | Kansas State University | Manhattan, KS, USA | LGU | TD, PR, NC | SAWAGEN, GE,<br>ILCI | POO, SM, TJF |
| PSU | Pennsylvania State University | University Park, PA,<br>USA | LGU | TD, PR | AM | JM, CMM, JRL |

<sup>a</sup> Institution type. NARI = National Agricultural Research Institute. AU = African University, NPRO = Non Profit Research Organization, LGU = Land Grant University. <sup>b</sup> Role in network: VD = Varietal Development, TD = Trait Development, PR = Pangenomic Resources, NC = Network Coordination. <sup>c</sup> Projects that supported the institutions participation in the network: SAWAGEN = "Improving sorghum adaptation in West Africa with genomics-enabled breeding" (2014–2019) and "Improving Sorghum Adaptation in West Africa with a Genomics-Enabled Breeding Network" (2019–2023), funded by USAID–Sorghum & Millet Innovation Lab. ILCI = "Innovation Lab for Crop Improvement" (2019–2025), funded by USAID. AVISA = "Accelerated Varietal Improvement and Seed Delivery of Legumes and Dryland Cereals in Africa" (2023–now) funded by Gates Foundation. AM = "Mining useful alleles for climate change adaptation from CGIAR gene banks" (2022–now) funded by Gates Foundation. GE = "Green Evolution - Accelerating Dryland Cereals Improvement for Africa" (2024–now) funded by Gates Foundation

**Extended Data Table 2:** Summary of phenotyping trials.

| Location name | Coordinates | *Striga*<br>infestation type | Trial type | Phenotypes collected <sup>a</sup> | Trial dates | Trial manager |
| --- | --- | --- | --- | --- | --- | --- |
| Niamey | 2°07'59.1"E<br>13°29'15.3"N | Managed stress | Pot (on-station, field soils) | S45, S60, S90, CBU, CBI, PBI | 2022, 2023, 2024 | INRAN breeding team |
| Konni | 5°28'88.20"E<br>13°81'79.31"N | Managed stress | Field station | PH, DF, HG, S45, S60, S90 | 2023, 2024 | INRAN breeding team |
| Konni | 5°16'31.7"E<br>13°50'07.4"N | Natural infested hotspot | Smallholder farm | PH, DF, HG, S45, S60, S90 | 2024 | INRAN breeding team |
| Zinder | 8°59'1"E<br>13°50'46"N | Uninfested control | Smallholder farm | PH, DF, HG, SD, PL | 2024 | University of Zinder |
| Niamey | 2°07'59.1"E<br>13°29'15.3"N | Uninfested control | Field station | PH, DF, HG, BM, PL, PW, SD, PL | 2024 | INRAN breeding team |
| Maradi | 13°48'N<br>07°05'E | Uninfested control | Field station | PH, DF, HG | 2024 | INRAN breeding team |
| Matameye | 8°49'47"E<br>13°42'51"N | Uninfested control | Smallholder farm | PH, DF, HG, SD, PL | 2024 | University of Zinder |

<sup>a</sup> Phenotypes: PH = plant height, HG = 100-grain weight, DF = Days to Flowering, BM = plant biomass, PL = Panicle length, PW = Panicle weight, SD = Stem Diameter, S45 = number of *Striga* plant 45 days after sowing, S60 = number of *Striga* plant 60 days after sowing, S90 = number of *Striga* plant 90 days after sowing. CBU = Crop biomass in uninfested control pot, CBI = Crop biomass in infested pot, PBI = Parasite above-ground biomass in infested pot, PBU = Parasite above-ground biomass in uninfested control pot.

